# Mitochondrial-Derived Compartments are Multilamellar Domains that Encase Membrane Cargo and Cytosol

**DOI:** 10.1101/2023.07.07.548169

**Authors:** Zachary N. Wilson, Matt West, Alyssa M. English, Greg Odorizzi, Adam L. Hughes

**Affiliations:** Department of Biochemistry, University of Utah School of Medicine, Salt Lake City, UT 84112, USA; Department of Molecular, Cellular, and Developmental Biology, University of Colorado Boulder, Boulder, CO 80309

## Abstract

Preserving the health of the mitochondrial network is critical to cell viability and longevity. To do so, mitochondria employ several membrane remodeling mechanisms, including the formation of mitochondrial-derived vesicles (MDVs) and compartments (MDCs) to selectively remove portions of the organelle. In contrast to well-characterized MDVs, the distinguishing features of MDC formation and composition remain unclear. Here we used electron tomography to observe that MDCs form as large, multilamellar domains that generate concentric spherical compartments emerging from mitochondrial tubules at ER-mitochondria contact sites. Time-lapse fluorescence microscopy of MDC biogenesis revealed that mitochondrial membrane extensions repeatedly elongate, coalesce, and invaginate to form these compartments that encase multiple layers of membrane. As such, MDCs strongly sequester portions of the outer mitochondrial membrane, securing membrane cargo into a protected domain, while also enclosing cytosolic material within the MDC lumen. Collectively, our results provide a model for MDC formation and describe key features that distinguish MDCs from other previously identified mitochondrial structures and cargo-sorting domains.

**SUMMARY:** Wilson and colleagues use electron tomography and time-lapse fluorescence microscopy to observe that mitochondrial-derived compartments (MDCs) are generated from outer mitochondrial membrane extensions that repeatedly elongate, coalesce, and invaginate to secure membrane cargo and cytosol within a distinct, protected domain.

## INTRODUCTION

Mitochondrial architecture is continuously remodeled to support the functional demands of the organelle and to preserve homeostasis. In actively growing cells, mitochondria form a dynamic, tubular network that is separated from the cytosol by two membranes. The outer mitochondrial membrane (OMM) creates an initial barrier and establishes connections with other organelles, while the inner mitochondrial membrane (IMM) creates an impervious barrier that protects the multitude of metabolic reactions occurring in the mitochondrial matrix (Pfanner et al., 2019; Harper et al., 2020). The IMM also dynamically invaginates to form cristae, which are imperative for efficient energy production and in the establishment of several distinct aqueous and membrane subdomains within mitochondria (Iovine et al., 2021). Because of the critical role mitochondria perform in cell metabolism, several investigations have analyzed the remodeling of mitochondria that occurs to match metabolic demand (Hackenbrock et al., 1966; Davies et al., 2012; Kondadi et al., 2020a; Kondadi et al., 2020b). Mitochondria also reorganize their architecture in response to diverse cellular stressors, and the formation of aberrant mitochondrial structures represent a hallmark phenotype of disease states and aging (Youle and van der Bliek, 2012; Hughes et al., 2012). Indeed, the failure to prune the mitochondrial network by removing impaired or damaged portions of the organelle can actively contribute to the progression of many neurodegenerative disorders (Palikaras et al., 2018; Killackey et al., 2020). Currently, the extent of remodeling mechanisms mitochondria use to respond to different stress conditions and to preserve mitochondrial health remains incompletely understood.

Various abiotic and biotic stressors can induce mitochondrial damage, leading to the separation and degradation of whole mitochondria through a variety of selective mitophagic processes (Killackey et al., 2020; Onishi et al., 2021). Mitophagy through the PINK1-Parkin pathway monitors mitochondrial health, in part by sensing deterioration of the mitochondrial membrane potential, which leads to the accumulation of PTEN-induced putative kinase 1 (PINK1) at the OMM. PINK1 subsequently recruits the E3 ligase Parkin, and together these proteins initiate a phosphorylation and ubiquitylation signaling cascade that marks mitochondria for mitophagic turnover (Lazarou et al., 2012; Kondapalli et al., 2012; Pickles et al., 2018). Conversely, receptor-mediated mitophagy uses distinct autophagic receptors localized on the mitochondrial surface to initiate mitophagy in response to diverse stress conditions, including starvation and hypoxia, or to remove mitochondria during development and cell differentiation (Schweers et al., 2007; Kanki and Klionsky, 2008; Zhang et al., 2008; Esteban-Martínez et al., 2017). In yeast, a reduction in metabolic demand and a switch to nitrogen starvation conditions leads to the expression and phosphorylation of the OMM-anchored mitophagy receptor Atg32 (Kanki et al., 2009; Okamoto et al., 2009; Aoki et al., 2011; Kanki et al., 2013), which in turn recruits autophagy machinery to initiate phagophore assembly and also recruits the yeast dynamin-related GTPase, Dnm1, so that portions of the mitochondrial network can be removed to facilitate mitochondrial turnover (Mao et al., 2013; Abeliovich et al., 2013).

Rather than reorganizing the entire mitochondrial network, some stress conditions induce mitochondria to sort cargo into distinct, membrane-bound domains leading to the piecemeal degradation of select mitochondrial cargo (Sugiura et al., 2014; Hughes et al., 2016). As a means for both steady-state turnover and in response to mild stress conditions, mitochondria form small vesicles (mitochondrial-derived vesicles, MDVs) approximately 50-160 nm in diameter that form by budding away from the mitochondrial network, encapsulating cargo from just the OMM or inclusive of both mitochondrial membranes and proteins from multiple mitochondrial subdomains (Soubannier et al., 2012a; Soubannier et al., 2012b; König et al., 2021). In response to mild oxidative stress, the formation of MDVs also involves the PINK1-Parkin pathway but occurs kinetically faster than full mitophagy (McClelland et al., 2016), and in some instances can compensate for the loss of mitophagy (Towers et al., 2021), all together suggesting that MDVs may act to preserve mitochondrial health prior to removal of whole mitochondria. Recently, a study that followed mitochondrial responses to *Toxoplasma gondii* infection observed that the targeting of the pathogen protein TgMAF1 to the OMM induced mitochondria to shed their OMM. However, rather than forming small OMM-derived MDVs, large (several microns in diameter) OMM-derived ring-shaped structures formed, called structures positive for the outer membrane (SPOTs), that robustly accumulated some OMM proteins while excluding intramitochondrial proteins (Li et al., 2022).

Previously, in the budding yeast, *Saccharomyces cerevisiae*, we identified a mitochondrial quality control pathway that also involves the selective sorting of mitochondrial proteins into a distinct domain called the mitochondrial-derived compartment (MDC). In old-aged yeast cells and in response to several acute stressors, mitochondria form large, spherical compartments that robustly sequester only a minor portion of the mitochondrial proteome (Hughes et al., 2016; Schuler et al., 2021). These MDCs are generated from a dynamic remodeling of mitochondrial membranes that rearrange at sites of contact with the ER and eventually form distinct spherical structures that contain resolvable lumens (English et al., 2020). Subsequently, MDCs are removed from mitochondria and delivered to yeast vacuoles for degradation, suggesting that MDCs act as a piecemeal autophagic mechanism that is induced to remodel or segregate select cargo from mitochondria (Hughes et al., 2016). Intriguingly, the primary cargo proteins identified within MDCs all include mitochondrial membrane proteins restricted to the OMM (Hughes et al., 2016; Wilson et al., 2023 *preprint*). However, the nature of MDC morphogenesis, the mechanisms involved in MDC formation, and the features of MDCs that distinguish them from other mitochondrial remodeling pathways all remains unresolved.

In this study, we used transmission electron microscopy and electron tomography to determine the ultrastructural morphogenesis of MDCs. We observed that MDCs form as large, multilamellar spherical compartments that frequently encase four membrane bilayers, all of which strongly labeled for the OMM protein Tom70. Using time-lapse fluorescence microscopy, we demonstrate that MDCs form through OMM extensions that repeatedly elongate, coalesce, and invaginate to create these compartments with layers of entrapped membrane. In doing so, MDCs engulf both OMM and cytosolic content, securing cargo into a distinct, protected domain, and thus provide evidence for how MDCs can robustly sequester certain cargo proteins from the OMM. Collectively, these results provide a model for MDC formation and define key features that distinguish MDCs from other previously identified mitochondrial structures and cargo-sorting domains.

## RESULTS

### Rapamycin treatment induces yeast to produce mitochondrial-derived multilamellar structures

Previously, we demonstrated that the inhibition of the mechanistic target of rapamycin (mTOR) robustly induces the formation of mitochondrial-derived compartments (MDCs) (Schuler et al., 2021). MDCs are novel mitochondrial subdomains characterized by their strong enrichment of a select portion of the mitochondrial proteome (Hughes et al., 2016). For example, in haploid yeast cells treated with rapamycin for two hours, we observe that ∼60% of cells form an MDC, demonstrated by the sequestration of the mitochondrial import receptor Tom70 into a large domain emerging from mitochondria that simultaneously excludes Tim50, an essential subunit of the Tim23 inner membrane translocase complex (Fig. 1, A and B, Hughes et al., 2016). In haploid yeast cells, MDCs typically resolve into large spherical domains, ∼400nm in mean diameter, that contain a resolvable lumen (Fig. 1, A and C). While these aspects of MDC formation have been previously characterized (English et al., 2020), the ultrastructural morphogenesis of MDCs remains unknown.

**Figure 1.**
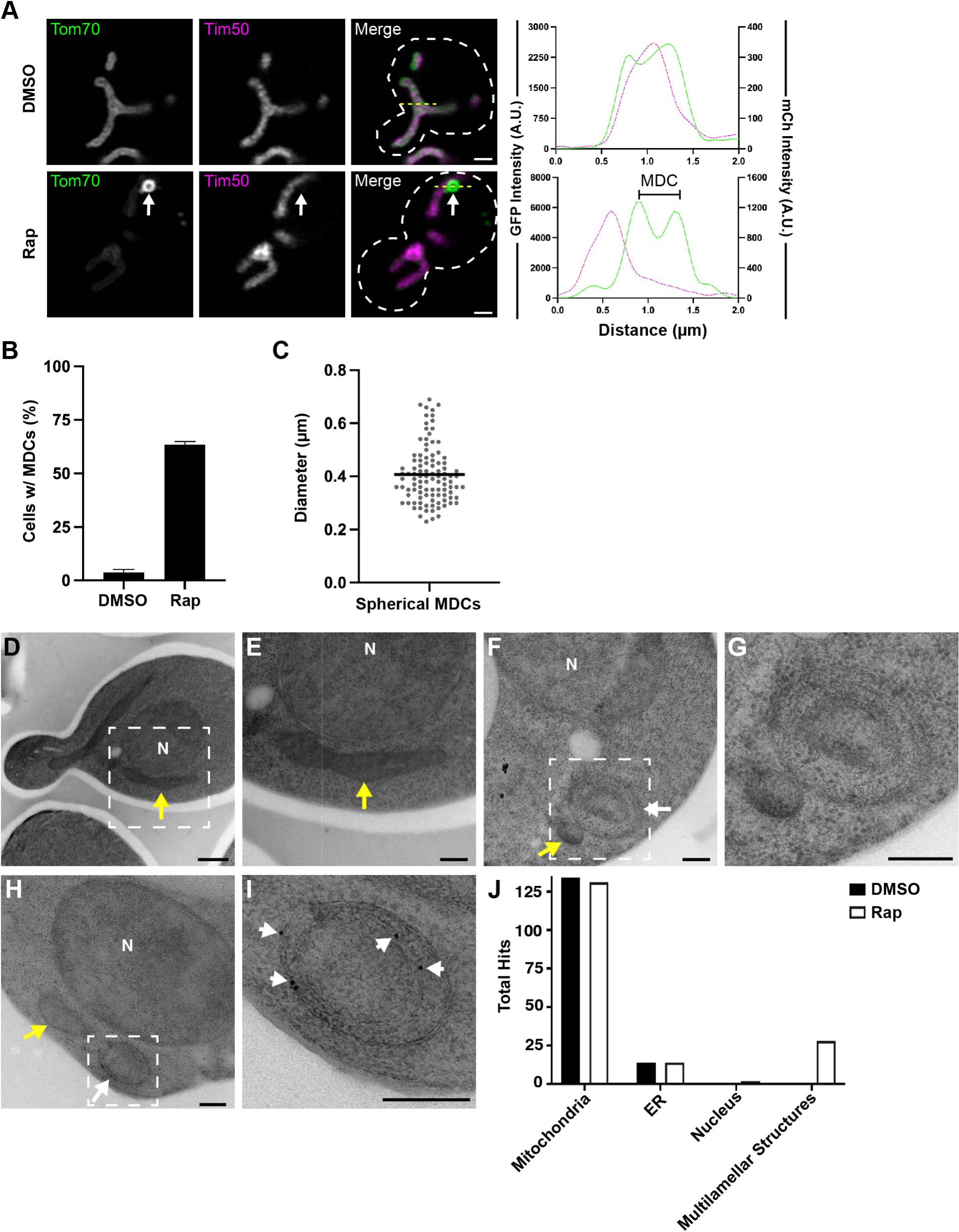
Rapamycin treated yeast produce mitochondrial-derived multilamellar structures. (A) Super-resolution confocal fluorescence microscopy images of DMSO or rapamycin (Rap)-treated haploid yeast cells expressing Tom70-yEGFP and Tim50-mCherry. MDCs are indicated by white arrows. Scale Bar= 1 µm. Yellow line marks the position of the line-scan fluorescence intensity profile shown on the right. Left and right Y axis correspond to Tom70-GFP and Tim50-mCherry fluorescence intensity, respectively. Bracket denotes MDC. (B) Quantification of MDC formation in DMSO or rapamycin treated yeast. Error bars show mean ± standard error of three replicates, *n* ≥ 100 cells per replicate. (C) Scatter plot showing the diameter of rapamycin-induced MDCs. Black line indicates the mean (0.41 µm) of *n* = 104 MDCs. (D-I) Thin-section TEM analysis of 80-nm cell sections from the same yeast strain analyzed above. Yeast were treated with either DMSO (D and E) or 200 nM rapamycin (F - I). White dotted-line squares in D, F, and H indicate the region magnified and shown in E, G, and I, respectively. Yellow arrows: mitochondria, white arrows: multilamellar structures, N: Nucleus. Scale Bars= (D) 500 nm (E-I) 200 nm. (H and I) Immunogold labeling with monoclonal antibodies targeting GFP and secondary antibodies conjugated to 10-nm gold particles. White arrowheads in I point to gold particles. (J) Quantification of the total anti-GFP immunogold particles that labeled the indicated cell structures from an analysis of >100 cell-sections.

To elucidate the structural characteristics of MDCs, we used thin-section transmission electron microscopy (TEM) to survey the ultrastructural morphogenesis of mitochondria in the same yeast strain that was analyzed in Fig. 1, A-C. Yeast were grown to log-phase, treated with DMSO (vehicle control) or 200 nM rapamycin for two hours, and then processed for TEM analyses by cryo-immobilization through high-pressure freezing followed by freeze-substituted fixation, a process that has been demonstrated to preserve membrane structure and limit fixation artifacts (West et al., 2011). In cell profiles from yeast treated with DMSO or rapamycin, mitochondria are readily observable as double-membrane bound organelles that form elongated tubules (longitudinal section) or small spherical organelles (cross section) with a darker luminal contrast compared to the yeast cytosol (yellow arrows, Fig. 1, D-F). Intriguingly, in cell profiles from yeast treated with rapamycin, we observed the formation of large (300-500 nm in diameter), spherical-shaped, multilamellar structures emerging or adjacent to mitochondrial tubules (Fig. 1, F and G). These mitochondrial-associated multilamellar structures were always observed in close proximity to or appeared directly attached to mitochondrial tubules. However, these structures appeared distinct from typical mitochondria because they contained multiple (>2) membrane bilayers and had a lighter luminal contrast (Fig. 1, F and G). These multilamellar structures appeared in ∼2% of cell sections surveyed (out of >800 cell profiles) near our expected frequency (∼3%) for capturing a putative MDC structure by thin-section electron microscopy. We confirmed that these structures were derived from mitochondria as they labeled specifically with antibodies conjugated to 10-nm colloidal gold particles that targeted Tom70-GFP (Fig. 1, H-J). The labeling specificity of these antibodies is demonstrated by the high frequency by which we observed gold particles at mitochondria compared to other membrane-rich organelles, such as the ER and nucleus (Fig. 1 J). Together, these results demonstrate that rapamycin treatment induces yeast to produce mitochondrial-derived multilamellar structures.

Initially, we characterized the formation of MDCs in aged yeast cells and in yeast treated with the Vacuolar H^+^-ATPase inhibitor, Concanamycin A (ConcA), which mimics the alkalinization of vacuoles that occurs during yeast aging (Hughes et al., 2016; Hughes et al., 2012). While treating cells with 500 nM ConcA induces MDC formation, MDCs are less frequent (forming in ∼40% of cells) and resolve into spherical domains that are slightly smaller (∼360nm in mean diameter) than those produced after rapamycin treatment (Fig. S1, A-C; Schuler et al., 2021). Despite this reduction in MDC formation upon ConcA treatment compared to rapamycin treatment, we also observed the formation of mitochondrial-associated multilamellar structures in cell profiles derived from yeast that had been treated with 500nM ConcA for two hours (Fig. S1 D). These multilamellar structures were also mitochondrial-derived as immunolabeling demonstrated that they contained Tom70-GFP (Fig. S1, D and E). Considering the mitochondrial network is preserved in rapamycin-treated cells, and that MDCs form more frequently upon rapamycin treatment, we focused the rest of our EM analyses on yeast treated with rapamycin.

### Mitochondrial-derived multilamellar structures enrich for Tom70-GFP and exclude Tim50-mCherry

A defining feature of MDCs is the exclusion of most mitochondrial proteins, including the inner membrane protein, Tim50 (Fig. 1 A). Notably, a dual-labeled immunoelectron analysis demonstrated that while the rapamycin induced mitochondrial-derived multilamellar structures labeled strongly for Tom70-GFP, they were largely devoid of Tim50-mCherry (Fig. 2). A serial-section reconstruction derived from thin-section TEM images of a dual immuno-labeled multilamellar structure revealed that this structure formed an elongated, spherical compartment that appeared to be enclosing at two tapered ends (Fig. 2, A-E; and Video 1). Other than the two tapered ends, this structure appeared completely enclosed. However, we cannot exclude the possibility that small (∼10-30nm) openings exist, and we did not capture the entirety of this mitochondrial-derived multilamellar structure (Fig. 2, A-E; and Video 1). The outer compartment was ∼520nm in diameter and was bound by two closely-apposed membrane bilayers that surrounded a second compartment (∼490nm in diameter) also formed by two closely-apposed membrane bilayers (Fig. 2, A-C; and Video 1). Additionally, this mitochondrial-derived multilamellar structure appeared to be directly adjacent to a mitochondria-ER contact site, as the ER was identified based on the size and contrast staining that were consistent with prior observations for ER membranes (Fig. 2, D and E, ER is colored in yellow; and Video 1; West et al., 2011), and near an additional double-membrane vesicular structure that labeled strongly for Tom70-GFP (Fig. 2, D and E, labeled in green; and Video 1).

**Figure 2.**
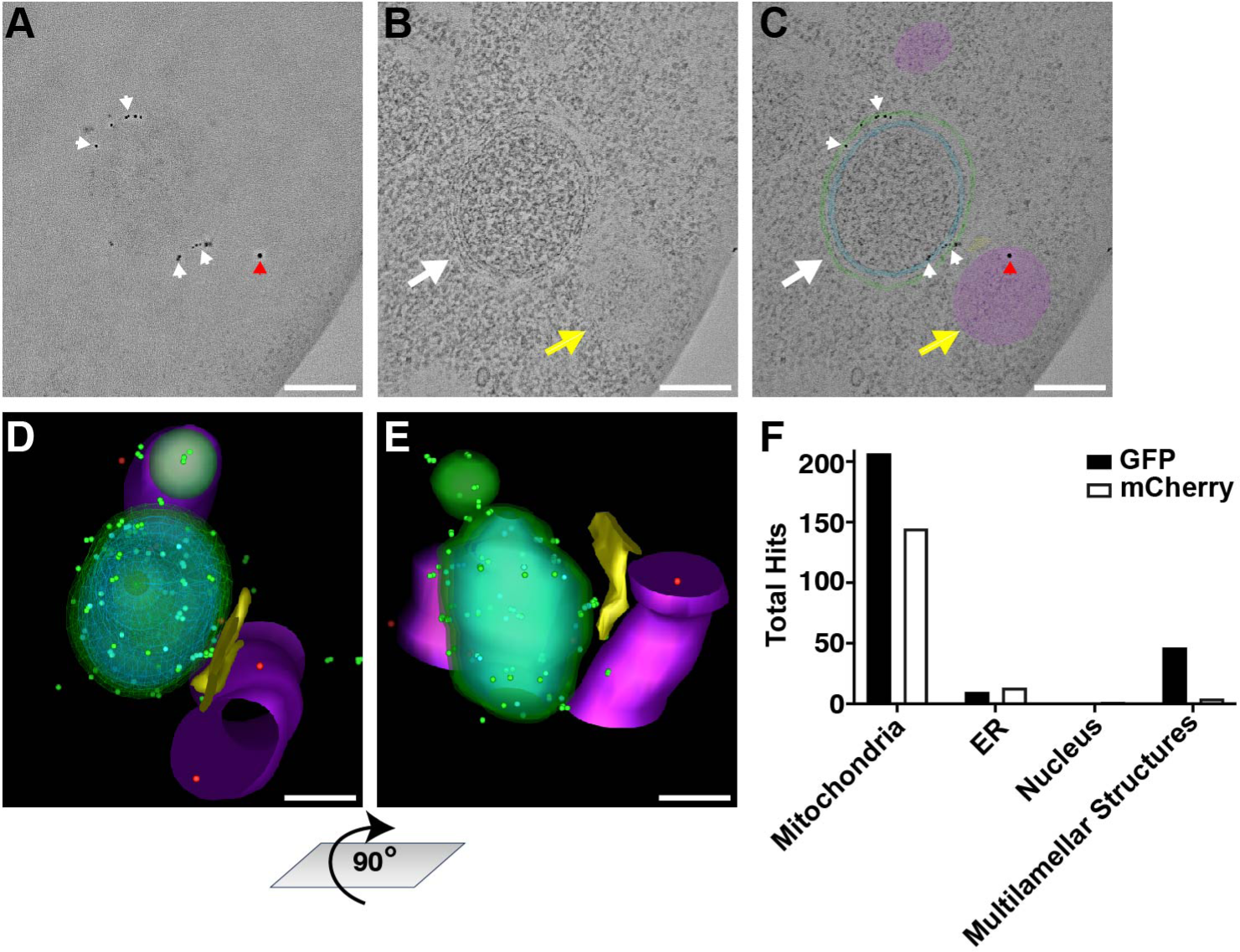
Mitochondrial-derived multilamellar structures are enriched for Tom70-GFP and exclude Tim50-mCherry. (A-E) Dual-immuno tomography obtained from five 90-nm cell sections of a yeast cell expressing Tom70-yEGFP and Tim50-mCherry treated with 200 nM rapamycin. Secondary antibodies conjugated to 6-nm or 10-nm gold particles targeted primary antibodies for GFP or mCherry, respectively. Scale Bar= 200 nm. See related Video 1. (A) Tomograph of the immune-reactive surface of the middle section (cell section #3). White arrowheads point to 6-nm gold particles, red arrowheads point to 10-nm gold particles. (B) Tomograph of the mid-plane of the middle section (cell section #3). Yellow arrow: mitochondria, white arrow: multilamellar structure. (C) Model overlay of section 3. Purple labels mitochondria, green labels the outer layer of doublet membranes, while cyan labels the internal layer of doublet membranes in the mitochondrial-derived multilamellar structure. (D) 3D model of the dual-immuno tomography described above. The mitochondrial-derived multilamellar structure is wire-modeled with the limiting membrane of the outer double-membrane labeled in green and the inner doublet membrane labeled in cyan. 6nm-gold particles appearing on the outer surface are overlayed with green spheres, while 6nm-gold particles appearing within the multilamellar structure are overlayed with cyan spheres. 10-nm gold particles are overlayed with red spheres. Mitochondria: purple, ER: yellow. (E) Same 3D model as shown in D rotated vertically 90°. (F) Quantification of the total anti-GFP 6-nm and anti-mCherry 10-nm immunogold conjugated secondary antibodies that labeled the indicated cell structures from a thin-section TEM analysis of 178 cell-sections from yeast expressing Tom70-yEGFP and Tim50-mCherry that were treated with 200 nM rapamycin.

Antibodies conjugated to 6-nm colloidal gold particles that targeted Tom70-GFP could be observed throughout the reconstructed multilamellar structure, including within the interior (cyan-labeled dots), between and on both sets of double membrane compartments (cyan-labeled dots), and externally on the surface of the larger compartment (green-labeled dots) (Fig. 2, A and C-E; and Video 1). Conversely, antibodies conjugated to 10-nm colloidal gold particles that targeted Tim50-mCherry were only observed along mitochondrial tubules and unenriched within the adjacent multilamellar compartment (Fig. 2 B-E; and Video 1). A larger dual-labeled immunoelectron analysis of 178 cell profiles revealed that antibodies detecting Tim50-mCherry are consistently absent from these mitochondrial-derived multilamellar structures (Fig. 2 F). The absence of Tim50-mCherry within these domains cannot be attributed simply to the lower protein abundance of Tim50 compared to Tom70, as antibodies against either protein strongly labeled mitochondria but Tim50 was more depleted in the multilamellar structures compared to the reduction in labeling observed at mitochondria (Fig. 2 F).

### Mitochondrial-derived multilamellar structures contain sets of paired membrane bilayers

Prompted by our observations from thin-section TEM, we examined the ultrastructure of mitochondria after rapamycin treatment by thick-section electron tomography. Electron tomography from yeast treated with rapamycin for two hours revealed the formation of spherical, multilamellar structures (labeled green) that were bound by two (Fig. 3 A-C; and Video 2) or four membrane bilayers (Fig. 3 D-F; and Video 3) in contact with mitochondrial tubules. All of these structures contained a lighter luminal contrast staining distinct from the contrast staining observed in the adjacent mitochondrial tubules and comparable to the surrounding cytosol (Fig. 3; and Videos 2 and 3). Measuring from the limiting membrane, the smaller double-membrane structures were ∼135 nm and ∼170 nm in diameter and were reminiscent of the smaller, double-membrane structure strongly labeled with Tom70-GFP seen previously (compare Fig. 3 A-C to Fig. 2 D and E). Also similar to the multilamellar compartment observed in Fig. 2, the larger, multilamellar structure was bound by a set of two closely-apposed membrane bilayers (∼420 nm in diameter) that surrounded an internal layer of closely-apposed paired membranes (∼370 nm in diameter). The diameters of these larger, multilamellar structures are consistent with our measurements of MDC diameter from super-resolution fluorescence microscopy (Fig. 1 C). Noticeably, each of these multilamellar structures were near an ER-mitochondria contact site or directly in contact with the ER, which was identifiable based on the continuity of the ER membranes that also contained areas with bound ribosomes (Fig. 2 and 3; and Videos 1-3; ER labeled in yellow). These observations are consistent with results from a prior investigation that showed that MDCs form at ER-mitochondria contact sites and also require ER-mitochondria contact sites for MDC biogenesis (English et al., 2020). Interestingly, another serial-section reconstruction of a large (∼720nm in diameter) multilamellar structure that robustly labeled with antibodies targeting Tom70-GFP showed extensive ER contact, potentially indicating that ER contact with MDCs increases as these structures grow in size (Fig. S2 A-C; and Video S1). Altogether, the results from our electron microscopy analyses support the interpretation that MDCs are formed from multiple layers of mitochondrial membrane and contain luminal content distinct from the mitochondrial matrix.

**Figure 3.**
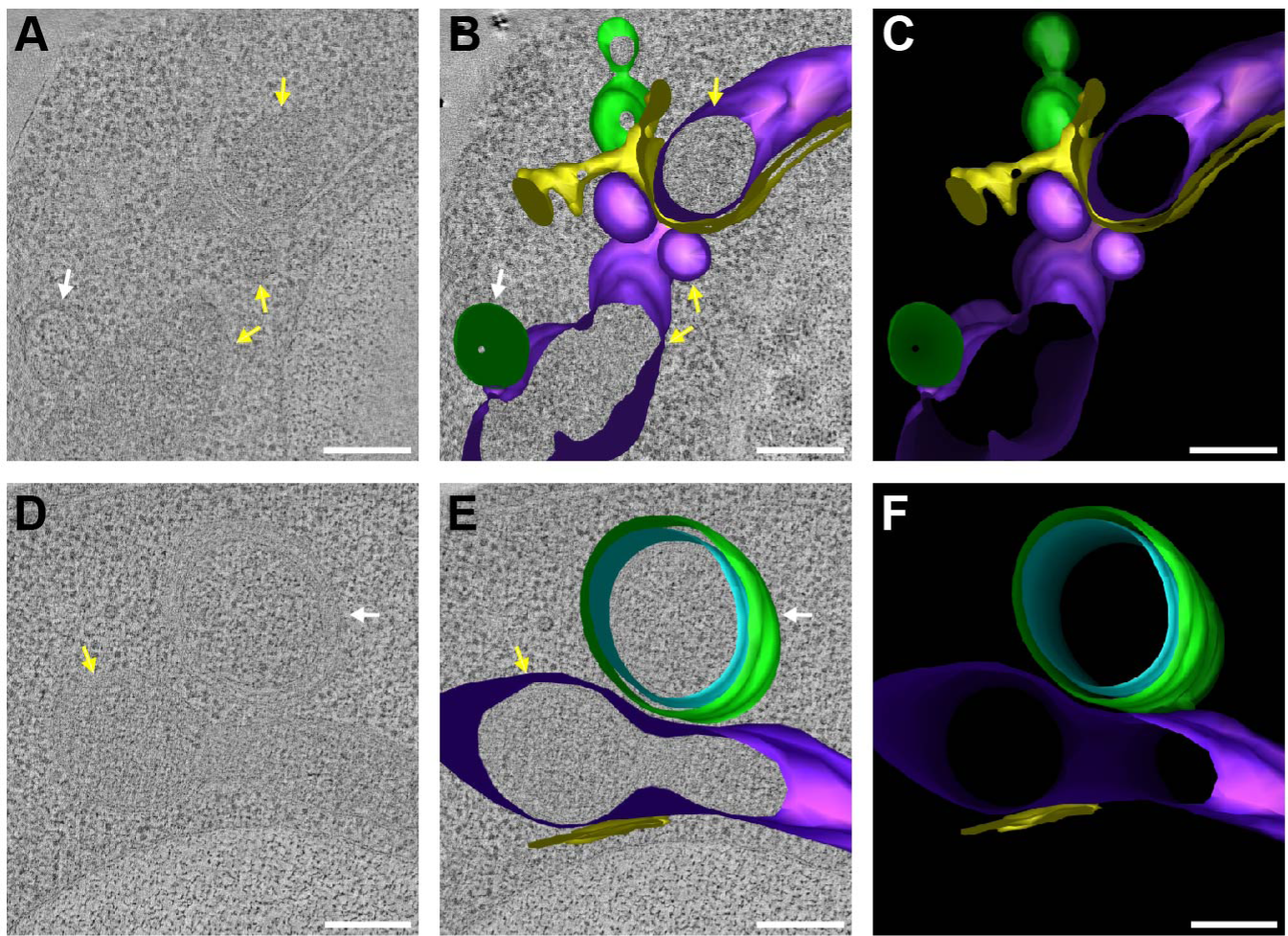
Mitochondrial-derived compartments contain sets of paired membranes. (A - F) 2D cross sections and 3D models derived from 200-nm cell sections of yeast cells treated with 200 nM rapamycin. Scale bar= 200 nm. See associated Videos 2 and 3. (A) Tomograph of a small, putative MDC bound by a single double-membrane. Yellow arrows: mitochondria, white arrow: MDC. (B) Model overlay of the tomograph shown in A. (C) 3D model of small, putative MDCs bound by a single double-membrane (labeled green). Mitochondria: purple, ER: yellow. See associated Video 2. (D) Tomograph of a larger, putative MDC bound by four membrane bilayers. Yellow arrow: mitochondria, white arrow: MDC. (E) Model overlay of the tomograph shown in D. (F) 3D model of the larger, putative MDC bound by two closely-apposed paired membranes. The limiting membrane of the outer doublet membrane is labeled green, while the internal doublet membrane is labeled cyan. Mitochondria: purple, ER: yellow. See associated Video 3.

While autophagosomes frequently associate with mitochondria, and it has been reported that mitochondrial membranes and ER-mitochondria contact sites support autophagosome biogenesis (Hailey et al., 2010; Hamasaki et al., 2013), we considered it unlikely that autophagosomal membranes were involved in generating the membrane-enriched mitochondrial-derived compartments we observed in our EM analyses. Previously, we demonstrated that both the core autophagy machinery and the yeast mitophagy receptor Atg32 are not required for MDC formation (Hughes et al., 2016). Furthermore, when we analyzed the localization of rapamycin-induced GFP-Atg8 foci compared to MDCs to assess if MDCs are bound by autophagosomal membranes, we observed that GFP-Atg8 foci did not co-localize with MDCs and were only within close proximity to MDCs about 15% of the time (orange arrows, Fig. S2, D-F). We also frequently observed MDCs near the cell periphery, while autophagosomes form and are often observed in close proximity to the vacuole (Fig. S2, G-J; Suzuki et al., 2013), and by electron tomography, autophagosomes often contained internal vesicles, which we have yet to see inside MDCs (Fig. S2, G-J, autophagosome labeled in orange). Moreover, our immunolabeling experiments for Tom70-GFP demonstrated that Tom70-GFP is found at both the surface and within internal membranes of MDCs, altogether indicating that MDCs are not additionally bound by autophagosomal membranes.

### Mitochondrial-derived compartments form through membrane extension intermediates

To investigate how mitochondria rearrange to form MDCs and capture the layers of membrane we observed in our ultrastructural analyses, we followed MDC biogenesis using time-lapse imaging of live yeast cells by fluorescence microscopy. Yeast expressing Tom70-GFP alone or Tom70-GFP and Tim50-mCherry were imaged every minute over a two-hour time course after MDC formation was induced via treatment with 200 nM rapamycin. Often within the first twenty minutes, we observed a membrane extension containing only Tom70-GFP that would extend along or emerge from mitochondria and subsequently fold back on itself to create a bright, spherical focus enriched for Tom70-GFP (Fig. 4 A-C; and Videos 4 and 5). Through seven separate experiments that captured 52 rapamycin-induced MDC biogenesis events, we observed that 43 of the MDC forming events (∼83%) began through a membrane extension intermediate that subsequently coalesced into a Tom70-GFP focus with greater fluorescent intensity than Tom70-GFP on the mitochondrial tubule (Fig. 4 B). Sometimes these bright Tom70-GFP foci would grow into large spherical domains with resolvable lumens that we have been defining as MDCs. However, frequently we found that the Tom70-GFP foci would repeat the process described above, continuing to grow and extend, creating bright, elongated extensions that invaginated prior to resolving into a large spherical compartment with a resolvable lumen (Fig. 4 C and Video 5; Fig. S3 A; and Video S2). These examples of Tom70-GFP positive membranes that appear to repeatedly extend and fold inward provide an explanation for how MDCs contain layers of membrane as observed by our ultrastructural analyses. This process of MDC formation also provides a potential explanation for how proteins trapped in MDCs become strongly enriched within these domains because they would be additionally captured within the internal membrane layers of MDCs.

**Figure 4.**
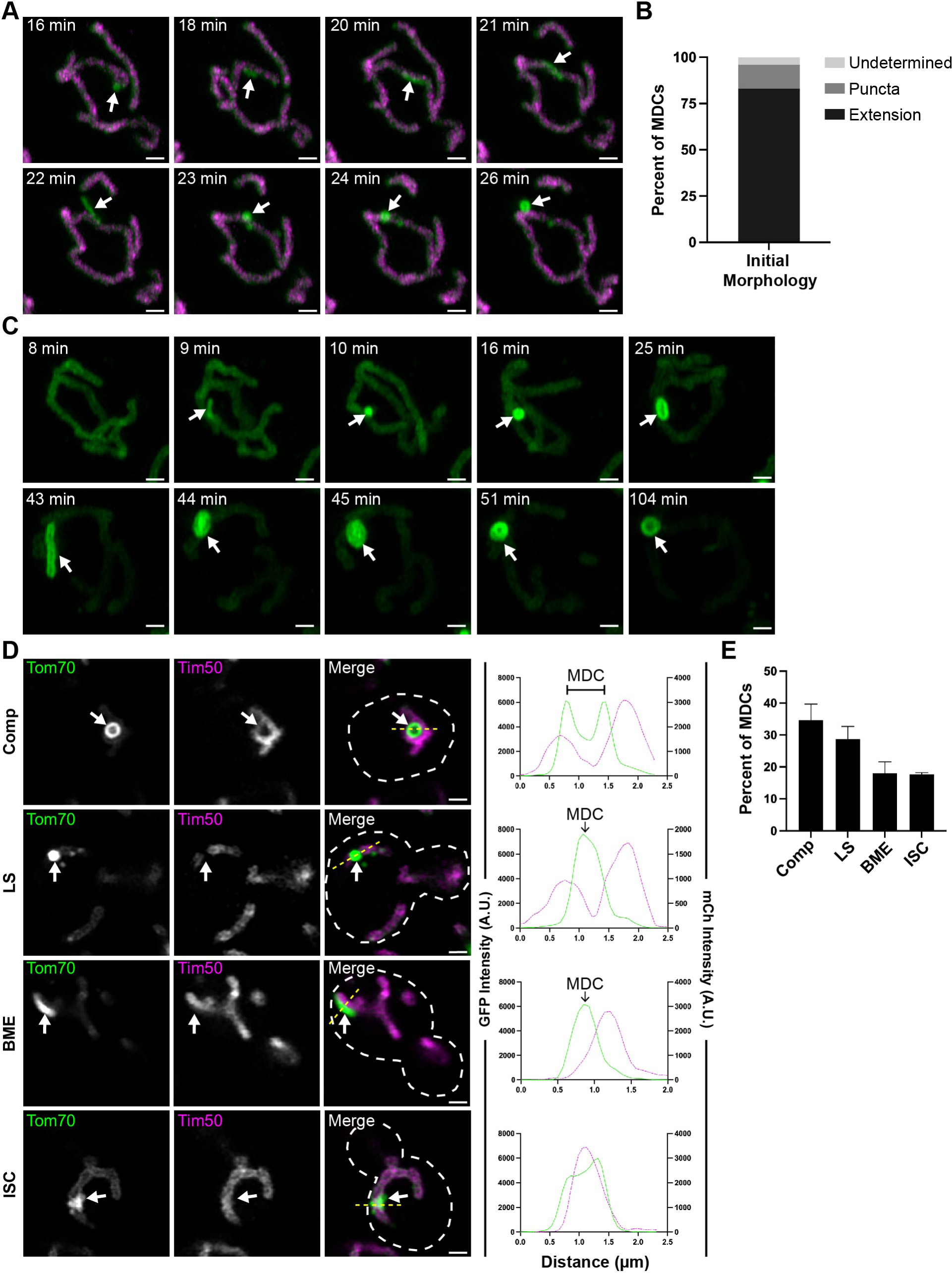
Mitochondrial-derived compartments form through membrane extension intermediates. (A) Super-resolution time-lapse images of rapamycin-induced MDC formation in yeast cells expressing Tom70-yEGFP and Tim50-mCherry. Images were acquired over 120 minutes (min). White arrows mark MDC. Scale bar = 1 µm. See associated Video 4. (B) Quantification of the initial morphology of Tom70-yEGFP structures during MDC biogenesis. *n*= 52 events from 7 experiments (C) Super-resolution time-lapse images of rapamycin-induced MDC formation in yeast cells expressing Tom70-yEGFP. Images were acquired over 120 minutes (min). White arrows mark MDC. Scale bar = 1 µm. See associated Video 5. (D) Representative super-resolution confocal fluorescence microscopy images of diverse MDC morphologies observed in rapamycin treated haploid yeast expressing Tom70-yEGFP and Tim50-mCherry. MDC structures are indicated by white arrows. Scale Bar= 1µm. Yellow line marks the position of the line-scan fluorescence intensity profile shown to the right. Left and right Y axis correspond to Tom70-yEGFP and Tim50-mCherry fluorescence intensity, respectively. (E) Quantification of the MDC morphologies shown in C as a percent of total MDCs. Error bars show mean ± standard error of three replicates, *n* > 60 MDCs per replicate.

Because our time-lapse imaging demonstrated that MDCs form through dynamic rearrangements of mitochondrial membranes that frequently but not always resolve into large spherical compartments, we quantitatively assessed the structural diversity of MDCs at the two-hour time point post rapamycin treatment. We binned the MDC morphologies that we observed into four categories: large spherical compartments with a resolvable lumen (compartment), bright spherical foci without a resolvable lumen (large spheres), bright membrane extensions (BME), or MDCs that formed as amorphous structures we defined as irregular-<u>s</u>haped Tom70-GFP-enriched <u>c</u>lusters (ISC) (Fig. 4 D). From this quantitative analysis we observed that most MDCs resolved into large spherical domains with (∼33%) or without (∼28%) resolvable lumens, consistent with our prior definition of MDCs (Fig. 4 E; Hughes et al., 2016; English et al., 2020). However, the final third of MDC structures were evenly split between BME and ISC morphologies, demonstrating that either MDCs continue to form over a long time period or that they do not always form clear compartment-like structures (Fig. 4 E). In support, we occasionally observed multiple MDCs forming in one cell (Fig. S3 A; and Video S2) and also captured multiple MDCs with diverse morphologies, all within one cell at the two-hour time point post rapamycin treatment (Fig. S3 B). While rare, we also observed MDCs with clear, resolvable, internal membrane invaginations (Fig. S3 C). These observations further supported our results that MDCs encapsulate layers of membrane and demonstrated that this feature of MDCs can be observed in a steady-state analysis of MDC morphology.

### Tom70-GFP-IAA7 is protected within MDCs from the auxin-degron system

Our time-lapse imaging experiments support our ultrastructural analyses that MDCs form as multilamellar compartments through repeated engulfment of the OMM. These results suggest that outer membrane proteins enclosed within the limiting membrane of MDCs should be protected from cytosolic degradation machinery. To assess this hypothesis experimentally, we fused an auxin-inducible degron to the C-terminus of Tom70-GFP (Tom70-GFP-IAA7; Nishimura et al., 2009). In cells treated with 1 mM indole-3-acetic acid (auxin), Tom70-GFP-IAA7 was rapidly degraded within the first thirty minutes (Fig. 5 A). While Tom70-GFP-IAA7 was rapidly degraded after auxin treatment, auxin had no detectable effect on the protein levels of several mitochondrial proteins, including the Tom70 paralog, Tom71, demonstrating that the auxin-induced degradation of Tom70-GFP-IAA7 is selective (Fig. 5 A). In diploid yeast cells expressing Tom70-GFP-IAA7 from one endogenous locus and Tom70-mCherry from the other locus, auxin treatment led to the near complete depletion of Tom70-GFP-IAA7 throughout the entire mitochondrial network while Tom70-mCherry remained unaffected (Fig. 5 B). Rapamycin treatment led to a robust sequestration of both Tom70-GFP-IAA7 and Tom70-mCherry into MDCs (Fig. 5 C, top panels), demonstrating that the auxin-inducible degron did not alter the recruitment of Tom70-GFP into MDCs. Intriguingly, when we treated with rapamycin for two-hours to establish MDCs prior to auxin addition, we observed a nearly complete depletion of Tom70-GFP-IAA7 throughout the mitochondrial tubule (Fig. 5 C, yellow arrows in Rap + Auxin panels), while Tom70-GFP-IAA7 remained protected within MDCs (Fig. 5 C, white arrows in Rap + Auxin panels). The protection of Tom70-GFP-IAA7 could also be observed in whole-cell lysates analyzed via western blot as Tom70-GFP-IAA7 was degraded at a slower rate in cells treated with rapamycin prior to auxin treatment compared to those treated with a vehicle control prior to auxin treatment (Fig. 5 D).

**Figure 5.**
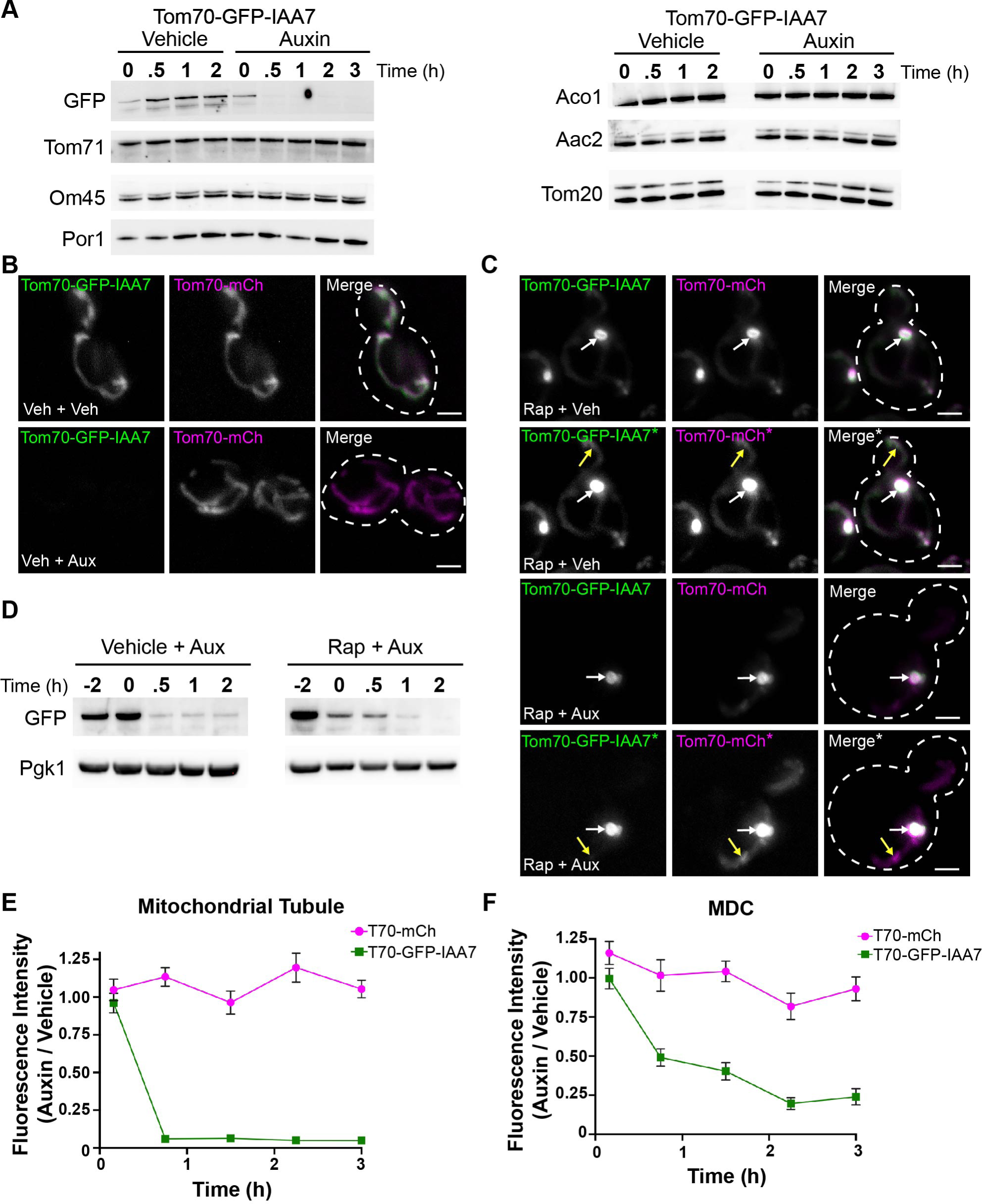
Tom70-GFP-IAA7 is protected within MDCs from the auxin-degron system. (A) Immunoblots of whole-cell protein extracts from yeast expressing Tom70-yEGFP-IAA7 and probed for the indicated mitochondrial proteins or GFP. Extracts were obtained at the indicated time points after treatment with either 1 mM auxin or an equivalent volume of 70% ethanol (Vehicle). (B) Max projections of widefield fluorescence microscopy images of yeast cells expressing Tom70-GFP-IAA7 and Tom70-mCherry. Images were taken 1.5 hours after treatment with either 1 mM auxin (Veh + Aux) or an equivalent volume of 70% ethanol (Veh + Veh), which were both added 2 hours after treatment with DMSO. Scale bar= 2 µm (C) Similar analysis as shown in B except yeast cells were treated with 200 nM rapamycin two hours prior to treatment with either auxin (Rap + Aux) or an equivalent volume of 70% ethanol (Rap + Veh). Yellow arrows indicate mitochondrial tubule, white arrows indicate MDCs. *Panels showing an increased pixel intensity. Scale bar= 2 µm. (D) Immunoblot of whole-cell protein extracts from yeast expressing Tom70-GFP-IAA7. Extracts were obtained at the indicated time points around treatment with 1 mM auxin. Auxin treatment occurred 2 hours after treatment with either 200 nM rapamycin or DMSO (Vehicle) and a whole-cell protein extract sample was obtained prior to drug treatment (−2). Pgk1 is provided as a loading control. (E) An analysis comparing the fluorescence intensities elicited by either Tom70-mCherry or Tom70-GFP-IAA7 within the mitochondrial tubule at the indicated time points. Fluorescence intensity is shown as a ratio of emissions from mitochondrial tubules of auxin-treated cells / vehicle-treated cells. (F) A similar analysis as shown in E except the fluorescence intensity ratios were derived from comparing emissions from MDCs of auxin-treated cells / vehicle-treated cells.

To further analyze the impact of MDC sequestration on the auxin-induced degradation of Tom70-GFP-IAA7, we performed a time-course experiment, capturing the fluorescence intensities of Tom70-GFP-IAA7 and Tom70-mCherry in cells that were either pre-induced or uninduced for MDC formation two-hours prior to treatment with either auxin or a vehicle control. Upon auxin or vehicle treatment, we captured live-cell images on large cell populations (n>100 cells) every 45 minutes for three hours. A plot providing the ratio of the mean fluorescence intensity of Tom70-GFP-IAA7 found within the mitochondrial tubule after auxin treatment compared to the vehicle control demonstrates the rapid and near complete removal of Tom70-GFP-IAA7 by the 45-minute time-point. Furthermore, the levels of Tom70-GFP-IAA7 in the mitochondrial tubule remained nearly undetectable throughout the entirety of the three-hour time-course experiment (Fig. 5 E). In comparison, the levels of Tom70-mCherry remained unaffected by auxin treatment highlighted by a fluorescence intensity ratio that consistently hovered around one (Fig. 5 E). In contrast to the rapid auxin-induced degradation of Tom70-GFP-IAA7 in mitochondrial tubules, Tom70-GFP-IAA7 sequestered in MDCs was protected from auxin-induced degradation, illustrated by the slower depletion of Tom70-GFP-IAA7 fluorescence intensity within MDCs (Fig. 5 F). By the end of the three-hour time course, the amount of Tom70-GFP-IAA7 observed within MDCs had diminished to a quarter of its initial fluorescence intensity but never dropped to the nearly undetectable levels caused by auxin-induced degradation within mitochondrial tubules (Fig. 5 F). While these results further demonstrate that Tom70-GFP-IAA7 is protected within MDCs, they also suggest that a significant portion of Tom70-GFP-IAA7 can still be degraded within MDCs or that Tom70-GFP-IAA7 within MDCs can escape back into mitochondrial tubules where it is subsequently degraded. Because the fluorescence intensity ratios plotted were derived from different cell populations captured over the time-course experiment, some of the reductions in Tom70-GFP-IAA7 levels observed in MDCs could also be attributed to the formation of MDCs that occur after Tom70-GFP-IAA7 has already been degraded.

### The MDC lumen contains cytoplasmic material

Next, we wanted to determine the nature of the material within the MDC lumen. Based on the appearance of the MDC lumen in our electron micrographs, we hypothesized that cytoplasm is engulfed within the MDC interior. We began by testing if cytoplasmic material is excluded from MDCs as it is from mitochondria. To do so, we expressed GFP in cells also expressing Tom70-mCherry and induced MDC formation with rapamycin treatment. In these cells, GFP filled the entire cytoplasm and nucleoplasm but was clearly excluded from the interior of yeast vacuoles and mitochondrial tubules (Fig. 6 A, yellow arrows). In contrast, we could distinguish neither an exclusion of cytoplasmic GFP from MDCs nor an enrichment of cytoplasmic GFP within MDCs (Fig. 6 A, white arrows), demonstrating that MDCs can be infiltrated with cytoplasmic material. Furthermore, as a comparison, we used an auxin-induced degron system to acutely dissolve the ER-mitochondria encounter structure (ERMES), which resulted in the appearance of swollen mitochondrial tubules and spheres as previously reported (John Peter et al., 2022). Even though mitochondrial architecture was lost in these cells, cytoplasmic GFP was still strongly excluded from the lumen of these aberrant mitochondria (Fig. 6 B, yellow arrows). Altogether, these results are consistent with our TEM analyses on MDC morphogenesis, as we observed a lighter luminal electron density within MDCs distinct from staining observed in adjacent mitochondria and comparable to the surrounding cytosol.

**Figure 6.**
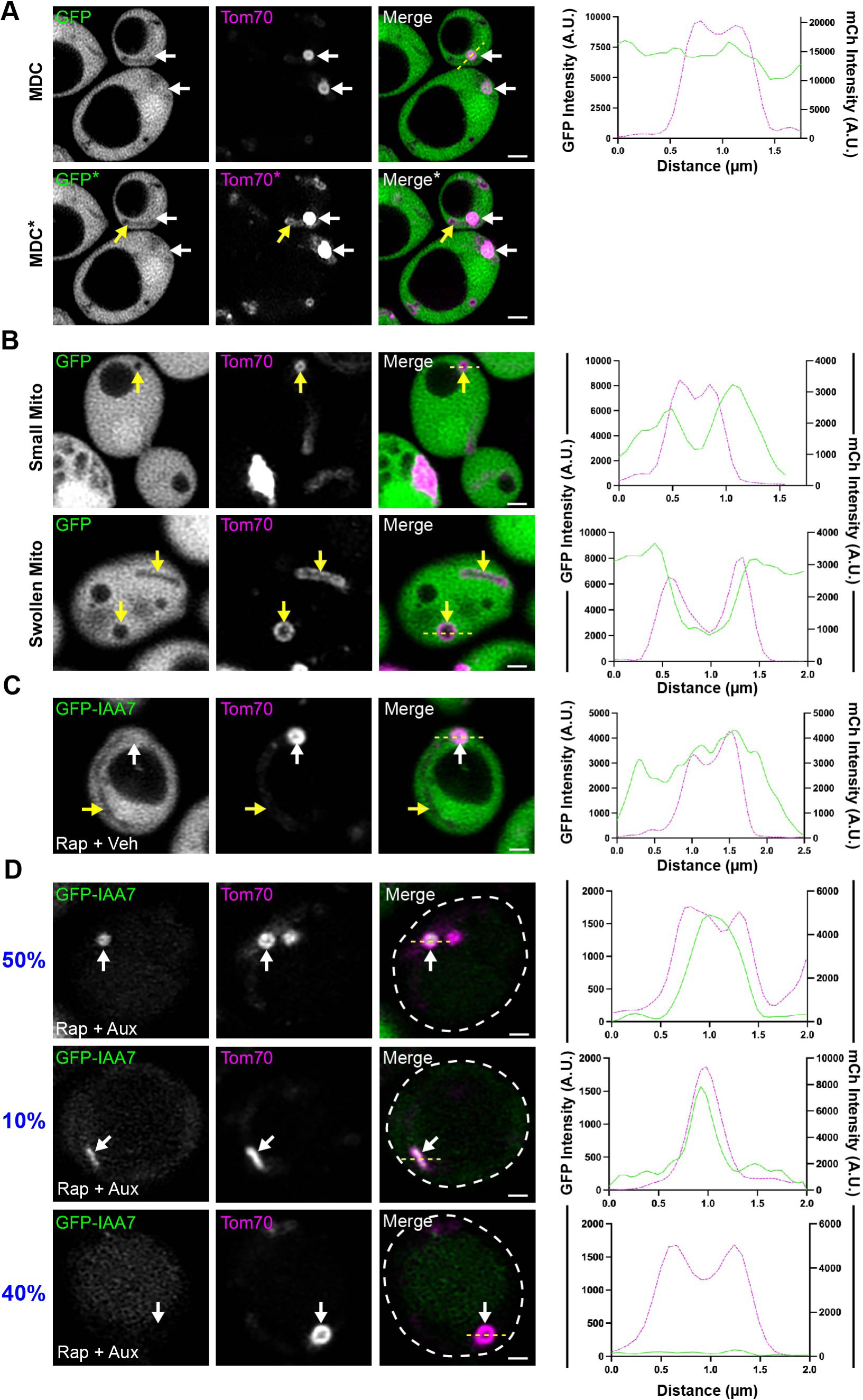
The MDC lumen contains cytoplasmic material. (A) Super-resolution confocal fluorescence microscopy images of rapamycin-induced MDC formation in yeast expressing yEGFP and Tom70-mCherry. MDCs are indicated by white arrows. Scale Bar= 1 µm. Yellow line marks the position of the line-scan fluorescence intensity profile shown to the right. Left and right Y axis correspond to yEGFP and Tom70-mCherry fluorescence intensity, respectively. *Panels showing an increased pixel intensity to observe Tom70-mCherry marked mitochondrial tubules indicated with a yellow arrow. (B) Super-resolution confocal fluorescence microscopy images of swollen mitochondria (yellow arrows) from yeast expressing yEGFP, Tom70-mCherry, Mdm12-AID-6xFLAG, and *Os*Tir1. Images were taken 3 hours after treatment with 1 mM auxin, which acutely swelled mitochondria through auxin-induced degradation of Mdm12-AID-6xFLAG. Scale Bar= 1 µm. Yellow line marks the position of the line-scan fluorescence intensity profile shown to the right. Left and right Y axis correspond to yEGFP and Tom70-mCherry fluorescence intensity, respectively. (C and D) Representative super-resolution confocal fluorescence microscopy images of yeast expressing yEGFP-IAA7 and Tom70-mCherry. After a two-hour treatment with 200 nM rapamycin, cells were subsequently treated with 70% ethanol (C; Rap + Veh) or 1 mM auxin (D; Rap + Auxin). Yellow arrows mark mitochondria while white arrows mark MDCs. Scale Bar= 1 µm. Yellow line denotes the position of the line-scan fluorescence intensity profile shown to the right. Left and right Y axis correspond to GFP and Tom70-mCherry fluorescence intensity, respectively. The blue percentages next to the panels shown in D indicate the frequency those results were observed from *n=*106 MDCs from four experiments.

While our results demonstrate that cytoplasmic GFP is not excluded from MDCs, it remained unclear if MDCs contained openings to the cytoplasm or if MDCs could fully entrap cytoplasmic material. To distinguish between these possibilities, we fused the auxin-induced degron to cytoplasmic GFP (GFP-IAA7), induced MDC formation, and subsequently treated cells with vehicle control or auxin. In vehicle-treated cells, we similarly observed that cytoplasmic GFP-IAA7 was not excluded from the interior of MDCs (Fig. 6 C). Strikingly, upon auxin treatment, we often observed that the only GFP-IAA7 signal that remained came from the interior of MDCs, demonstrating that MDCs could encase cytoplasmic GFP-IAA7 and protect it from auxin-induced degradation (Fig. 6 D, white arrow). We observed that cytoplasmic GFP-IAA7 was protected in both the large spherical MDCs (40% of MDCs) and in the bright, membrane extensions (10% of MDCs, Fig. 6 D) demonstrating that these bright extensions are elongated compartments with captured cytoplasmic GFP-IAA7. Importantly, we also found that in ∼50% of cases, MDCs could not protect cytoplasmic GFP-IAA7 from degradation (Fig. 6 D, bottom panels), suggesting that either these MDCs formed after GFP-IAA7 was degraded, or that in some instances, openings exist to allow exchange with the cytoplasm. Collectively, these results demonstrate that MDCs are a distinct reorganization of OMM, capable of entrapping layers of OMM and cytoplasmic content.

### The mitochondrial fission and fusion machinery perform competing roles in MDC formation

Our results suggest that MDCs form through the repeated elongation and closure of OMM-derived membrane extensions, suggesting that membrane remodeling machinery is involved in MDC biogenesis. To test whether the mitochondrial fission and fusion machinery is involved in MDC formation, we began by determining the localization of the mitochondrial fission and fusion GTPases, Dnm1 and Fzo1, respectively, compared to MDCs (Fig. 7 A-C). Consistent with previous observations, we observed that GFP-Dnm1 remains punctate on mitochondrial tubules but also strongly associates with the majority of MDCs upon MDC induction (Fig. 7, A and C; Hughes et al., 2016). Conversely, GFP-Fzo1 becomes robustly incorporated in 98% of observed MDCs and is present throughout the MDC structure (Fig. 7, B and C), providing evidence that the mitochondrial fusion machinery could be involved in forming MDCs. Because removal of Fzo1 (*fzo1*Δ) generates hyper-fragmented mitochondria (Hermann et al., 1998), we assessed the requirement of Fzo1 in MDC formation by analyzing MDC biogenesis in strains lacking *DNM1*, *dnm1*Δ and *dnm1*Δ*fzo1*Δ yeast, which maintains a tubular mitochondrial morphology. Surprisingly, *dnm1*Δ*fzo1*Δ yeast still robustly formed MDCs in response to both ConcA and rapamycin treatment (Fig. 7 D) and to a similar extent as was observed in wild-type and *dnm1*Δ cells. The continued formation of MDCs in *dnm1*Δ yeast matched our previous observations that the mitochondrial fission machinery is not required for MDC formation (Hughes et al., 2016). Furthermore, in assessing the structural diversity of MDCs in *dnm1*Δ cells we observed more MDCs resolving into large spherical domains and compartments compared to wild-type yeast (Fig. 7 E), indicating that Dnm1 may antagonize MDC formation by constricting or severing OMM proliferations before they round into spherical compartments. In *dnm1*Δ*fzo1*Δ cells, the structural diversity of MDCs resembled what we observed in wild-type cells, except that there was greater proportion of MDCs that formed bright membrane extensions (Fig. 7 E). While these results demonstrate that Fzo1 is not strictly required for MDC formation, they also implied that the role the mitochondrial fusion machinery performs in MDC formation might be masked by the complete absence of Dnm1 activity. Thus, we also assessed MDC formation in wild-type, *fzo1*Δ, and *dnm1*Δ*fzo1*Δ yeast that all ectopically expressed a temperature-sensitive version of Fzo1 (*fzo1-1*; Hermann et al., 1998). Notably, MDC formation was strongly impaired in *fzo1*Δ p*fzo1-1* cells after an acute one-hour shift to the non-permissive temperature of 37°C but not at the permissive temperature of 30°C (Fig. 7, F-H). Importantly, MDC formation still occurred in wild-type yeast expressing *fzo1-1* at 37°C, although we noted that MDC formation was consistently reduced at the higher temperatures overall because fewer MDCs also formed in control cells ectopically expressing wild-type Fzo1 (p*FZO1*; Fig. 7, F-H). Rather than forming MDCs, we frequently observed puncta containing only Tom70-GFP throughout the cell in *fzo1*Δ p*fzo1-1* cells treated with rapamycin at 37°C (Fig. 7 G, yellow arrows). These Tom70-GFP puncta did not appear enriched for Tom70-GFP as they were the same fluorescent intensity as that observed for the Tom70-GFP that remained in the fragmented mitochondria. Similar to our observations of MDC formation in *dnm1*Δ*fzo1*Δ cells, MDC formation was restored in *dnm1*Δ*fzo1*Δ pfzo1-1 cells at 37°C (Fig. 7 H). Thus, the mitochondrial fusion machinery is not absolutely required for MDC formation, but performs a distinct role counter-acting the activity of the mitochondrial fission machinery to allow MDCs to form from OMM proliferations.

**Figure 7.**
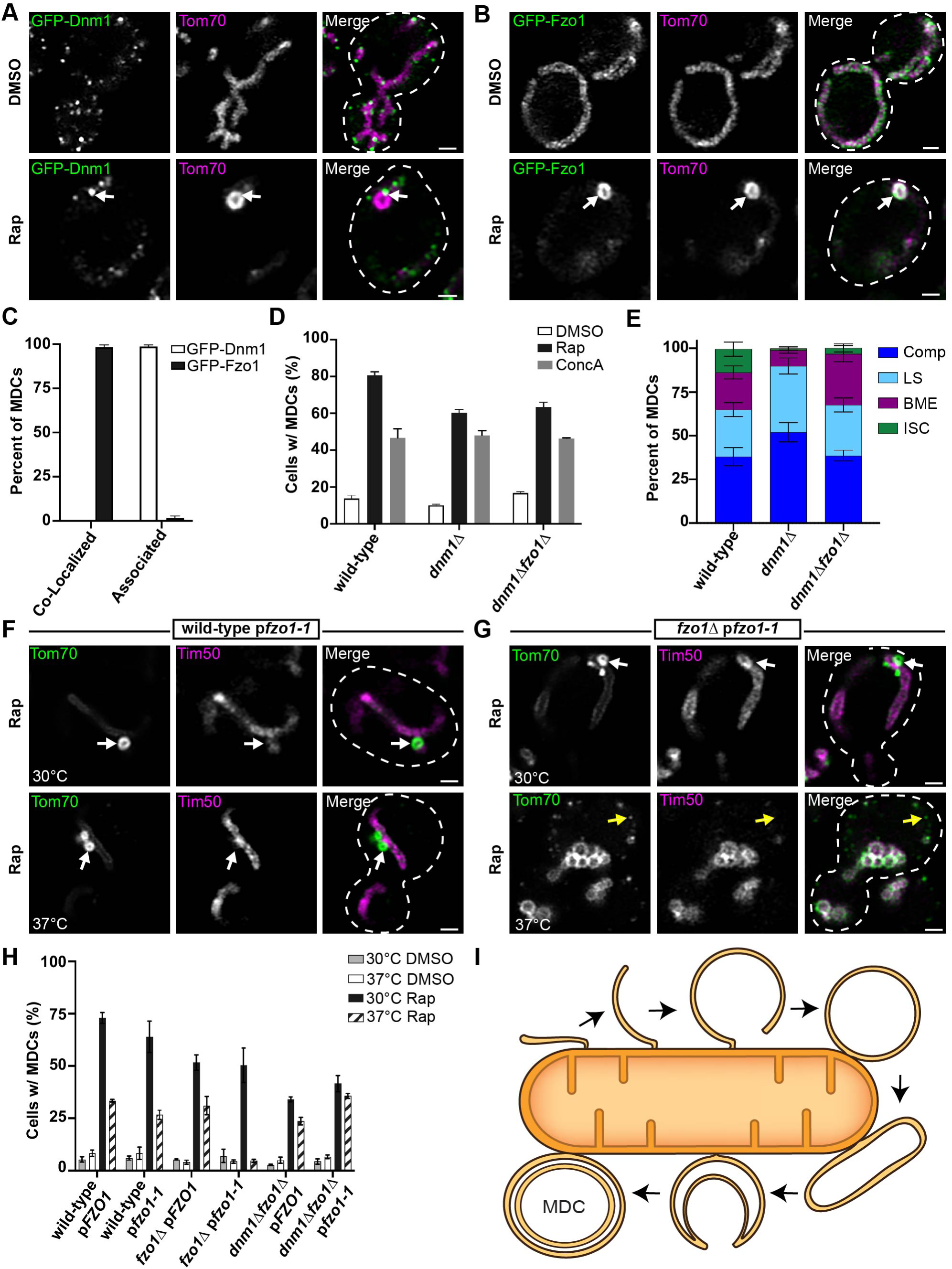
The mitochondrial fission and fusion machinery perform competing roles in MDC formation. (A) Super-resolution confocal fluorescence microscopy images of yeast expressing sfGFP-Dnm1 and Tom70-mCherry treated with either DMSO or 200 nM rapamycin (Rap). MDCs are indicated by white arrows. Scale Bar= 1 µm. (B) Super-resolution confocal fluorescence microscopy images of yeast expressing sfGFP-Fzo1 and Tom70-mCherry treated with either DMSO or 200 nM rapamycin (Rap). MDCs are indicated by white arrows. Scale Bar= 1 µm. (C) Quantification of the frequency sfGFP-Dnm1 foci or sfGFP-Fzo1 were co-localized with or closely associated to Tom70-mCherry-marked MDCs. Error bars show mean ± standard error of three replicates, *n* ≥ 100 cells per replicate. (D) Quantification of MDC formation in the indicated yeast strains upon treatment with either DMSO, concanamycin A (ConcA), or rapamycin (Rap). Error bars show mean ± standard error of three replicates, *n* ≥ 100 cells per replicate. (E) Quantification of the MDC morphologies observed in the indicated yeast strains after rapamycin treatment shown as a percent of total MDCs. Error bars show mean ± standard error of three replicates, *n* ≥ 100 cells per replicate. (F and G) Super-resolution confocal fluorescence microscopy images of wild-type (F) or *fzo1*Δ (G) yeast cells expressing *fzo1-1*, Tom70-yEGFP, and Tim50-mCherry treated with 200 nM rapamycin (Rap) at the indicated temperatures. MDCs are indicated by white arrows. Scale Bar= 1 µm. (H) Quantification of MDC formation in the indicated yeast strains upon treatment with either DMSO or rapamycin at the indicated temperatures. Error bars show mean ± standard error of three replicates, *n* ≥ 100 cells per replicate. (I) Model of MDC biogenesis from an OMM extension that forms a double-membrane compartment, elongating, and invaginating to form a multilamellar MDC.

## DISCUSSION

Mitochondria structurally reorganize to meet metabolic demands and, in times of stress, to preserve organelle homeostasis. Defects in mitochondrial dynamics and the observance of aberrant mitochondrial structures are hallmarks of disease states and a phenotype of aging cells (Youle and van der Bliek, 2012; Hughes et al., 2012). Previously, we identified a new structural domain of mitochondria, the MDC, that forms in aged yeast cells and mammalian cell culture in response to several acute stressors (Hughes et al., 2016; Schuler et al., 2020 *preprint*; Schuler et al., 2021). Here we used transmission electron microscopy and electron tomography to determine the ultrastructure of MDCs. We observed that MDCs form as large, multilamellar spherical compartments that frequently encase four membrane bilayers, whereby two closely-apposed membrane bilayers form an internal compartment that is surrounded by a second layer of closely-apposed double membrane bilayers. We demonstrate that MDCs form through OMM extensions that repeatedly elongate, coalesce, and engulf part of itself to create these compartments with layers of entrapped membrane. In doing so, MDCs engulf both OMM and cytosolic content, securing cargo into a distinct, protected domain. Overall, these results provide evidence of key features that distinguish MDCs from other previously identified mitochondrial structures and cargo-sorting domains.

Mitochondria form an elaborate architecture constructed of two membranes that are organized into several subdomains. While the OMM establishes the limiting membrane of mitochondria, the IMM surrounds the mitochondrial matrix and invaginates to create cristae, establishing several distinct aqueous and membrane subdomains (Iovine et al., 2021). Several membrane components and protein complexes embedded in the IMM are required to establish IMM architecture (Klecker and Westermann, 2021). Notably, the sharp-angled dimerization and oligomerization of F_1_F_O_-ATP synthase complexes within the IMM facilitates the generation and maintenance of mitochondrial cristae (Paumard et al., 2002; Davies et al., 2012). In the absence of ATP synthase dimerization, the IMM forms membrane sheets traversing the mitochondrial matrix, creating “onion-like” mitochondria as observed by electron microscopy (Paumard et al., 2002; Giraud et al., 2002; Davies et al., 2012). These mitochondria can form swollen spheres that are still delimited by the OMM (Paumard et al., 2002) but can appear to encase cytosol if they form depressed cup-like structures (Klecker and Westermann, 2021). As observed by EM, mitochondria can also form “onion-like” structures at sites of curved IMM septa (Harner et al., 2016) and in the absence of key organizational complexes, including the mitochondrial contact site and cristae organizing system (MICOS) and ERMES complexes (Stephan et al., 2021; Hobbs et al., 2001). Importantly, the “onion-like” structures observed in all of these scenarios with alterations of IMM architecture are still surrounded by a single OMM, contain layers of IMM, and still encase mitochondrial matrix. In contrast, the multilamellar compartments we observed forming upon MDC induction contain layers of OMM and are clearly capable of engulfing cytoplasmic content. Moreover, our previous analyses and observations within an accompanying manuscript all demonstrate that MDCs exclude content from the mitochondrial matrix, the IMM, and the intermembrane space (Hughes et al., 2016; Wilson et al., 2023 *preprint*), which is consistent with our observations of MDC ultrastructure and formation. Thus, while MDCs appear similar to mitochondria with aberrant IMM organization, they are actually a distinct remodeling of the OMM.

Our observations provide a model for how MDCs form through a membrane proliferation that is derived entirely from the OMM (Fig. 7 I). In this model, the generation of MDCs begins with an OMM extension that eventually rounds up and connects to create an initial double-membrane compartment that has engulfed cytosol. This compartment can continue to grow, elongate, and inwardly engulf part of itself and cytosol to create a compartment with layers of OMM encased in a concentric spherical compartment, where each compartment is bound by two closely-apposed membrane bilayers. In considering this model for MDC biogenesis, it was surprising to discover that Fzo1 is not strictly required for MDC formation because we predicted that Fzo1 activity may be necessary to either bring the membranes together or drive fusion. This result may indicate that the OMM proliferations do not fuse but instead form tight connections that we could not clearly resolve in our EM analyses. Alternatively, the topology of an OMM extension that encloses and subsequently forms an invaginated internal compartment may require a membrane scission event rather than membrane fusion (Zhen et al., 2021). It is also conceivable that MDCs form from an OMM extension that repeatedly folds back on itself to create karmellae on the mitochondrial surface. The formation of karmellae is a feature of membrane proliferations derived from the ER during ER microautophagy, which also appear as bright membrane extensions when visualized via fluorescence microscopy (Koning et al., 1996; Schäfer et al., 2019), and these ER-derived karmellae can round into large, spherical, multilamellar whorls (Wright et al., 1987; Schuck et al., 2014). Notably, we have not identified mitochondrial karmellae in our ultrastructural analyses and we also observed that the bright membrane extensions that appear during MDC biogenesis are capable of capturing cytoplasmic content (Fig. 6 D), suggesting that elongated compartments. However, we cannot fully exclude the possibility that MDCs form through multiple methods, including OMM-derived karmellae, which may be resolved with future studies that investigate the molecular mechanisms of MDC formation.

There are notable similarities between the MDC pathway and ER microautophagy. ER microautophagy can be induced by the overexpression of some resident ER membrane proteins, by ER stress, or by the aberrant retention of membrane proteins within the organelle (Wright et al., 1988; Schuck et al., 2009, Schäfer et al., 2020). The accumulation of these membrane proteins induces a dramatic proliferation of ER membranes that stack together as paired membrane doublets around the nucleus or in the cell periphery and also round into multilamellar whorls (Wright et al., 1988; Koning et al., 1996; Schuck et al., 2014). Subsequently, this distinct ER domain is degraded in vacuoles/lysosomes in a manner that does not rely on the core autophagy machinery or ER-specific autophagy receptors (Schuck et al., 2014). Prompted by these similarities, we observed in an accompanying manuscript that MDCs are also induced by the overexpression of many OMM proteins, and by the mistargeting of tail-anchored proteins to the OMM (Wilson et al., 2023 *preprint*). Here we show that MDCs, similar to ER microautophagy, form from a mitochondrial OMM proliferation to create a distinct membrane domain formed by closely-apposed, paired membrane bilayers, indicating that mitochondria can also generate dramatic membrane proliferations as a means to facilitate piecemeal autophagic turnover of the organelle.

Extensions, protrusions, and vesicles derived from the OMM have been observed in multiple cell types under both steady-state and pathological conditions (Soubannier et al., 2012a; Yao et al., 2021; Yamashita et al., 2016). Under mild stress conditions, mitochondria can release small vesicles that contain only the OMM or both mitochondrial membranes, delivering damaged protein cargoes, including intramitochondrial protein cargoes, to lysosomes for degradation (Soubannier et al., 2012a). Thus far, we have considered mitochondrial-derived compartments (MDCs) to be distinct from MDVs based on their size, mechanism of formation, and cargo proteins sequestered. The observations presented here further support that distinction, highlighted by our results showing that MDCs are OMM-enriched multilamellar compartments that engulf both OMM and cytosol. It seems possible that MDCs may provide cells a mechanism to sequester portions of the OMM or other cellular content that cannot be achieved by creating MDVs that are still delimited by the OMM. Interestingly, a recent study demonstrated that upon *Toxoplasma gondii* infection, the targeting of the pathogen protein *Tg*MAF1 to the OMM induced mitochondria to shed their outer membrane, creating large (several microns in diameter) ring-shaped structures, called SPOTs. These SPOTs robustly incorporated some OMM membrane proteins, while excluding intramitochondrial proteins, and included internal invaginations that were also morphologically reminiscent of ER whorls (Li et al., 2022). While it is currently unclear how SPOTs form, it would be interesting if they also form by invaginating OMM extensions and engulfing cytoplasmic material, similar to what we observed for MDC biogenesis. Altogether, it is clear that the remodeling of the OMM is a key mechanism by which cells preserve mitochondrial homeostasis. Understanding the heterogenous molecular mechanisms of this remodeling may provide new avenues to target pathological states that disrupt mitochondrial architecture.

## EXPERIMENTAL MODEL AND SUBJECT DETAILS

### Yeast Strains and Plasmids

All yeast strains are derivatives of *Saccharomyces cerevisiae* S288C (BY) (Brachmann et al., 1998) and are listed in Table S1. Deletion strains were created by one-step PCR-mediated gene replacement using the previously described pRS series of vectors (Brachmann et al., 1998; Sikorski and Hieter, 1989) and oligo pairs listed in Table S2. Correct gene deletions were confirmed by colony PCR across the chromosomal insertion site. Strains expressing proteins with attached C-terminal fluorescent proteins were created by one step PCR-mediated C-terminal endogenous epitope tagging using standard techniques and oligo pairs listed in Table S2. Plasmid templates for fluorescent epitope tagging were from the pKT series of vectors (Sheff and Thorn, 2004). Strains containing auxin-inducible degrons were constructed as described in the Auxin-Induced Protein Degradation section below. For all strains, correct integrations were confirmed by a combination of colony PCR across the chromosomal insertion site and correctly localized expression of the fluorophore by microscopy. Strains expressing proteins with attached N-terminal fluorescent proteins were derived from the SWAp-Tag library described in Weill et al., 2018 and were a gift from Maya Schuldiner. The plasmids used in this study are list in Table S3.

## METHOD DETAILS

### Yeast Cell Culture and Growth Assays

Yeast cells were grown exponentially for 15-16 hours at 30°C to a final optical (wavelength 600nm) density of 0.5 - 1 before the start of all experiments. This period of overnight log-phase growth was carried out to ensure vacuolar and mitochondrial uniformity across the cell population and is essential for consistent MDC formation. Unless otherwise indicated, cells were cultured in YPAD medium (1% yeast extract, 2% peptone, 0.005% adenine, 2% glucose). Otherwise, cells were cultured in a synthetic defined (SD) medium that contained the following unless specific nutrients were removed to select for growth or plasmid retention: 0.67% yeast nitrogen base without amino acids, 2% glucose, supplemented nutrients 0.072 g/L each adenine, alanine, arginine, asparagine, aspartic acid, cysteine, glutamic acid, glutamine, glycine, histidine, myo-inositol, isoleucine, lysine, methionine, phenylalanine, proline, serine, threonine, tryptophan, tyrosine, uracil, valine, 0.369g/L leucine, and 0.007 g/L para-aminobenzoic acid. Unless otherwise indicated, rapamycin, concanamycin A, and indole-3-acetic acid (auxin) were added to cultures at final concentrations of 200 nM, 500 nM, and 1 mM, respectively.

### MDC Assays

For MDC assays, overnight log-phase cell cultures were grown in the presence of dimethyl sulfoxide (DMSO) or the indicated drug for two hours. For MDC assays with cells containing plasmids, overnight log-phase yeast cultures grown in selective SD medium were back-diluted to an OD_600_=0.1-0.2 in YPAD medium and allowed to grow for at least 4 hours prior to MDC induction. For the temperature-sensitive MDC assays, cultures were shifted to the indicated temperatures 1 hour prior to MDC induction. Prior to visualization, cells were harvested by centrifugation, washed once, and resuspended in 100mM HEPES containing 5% glucose. Subsequently, yeast were directly plated onto a slide at small volumes to allow the formation of a monolayer and optical z-sections of live yeast cells were acquired with a ZEISS Axio Imager M2 or for super-resolution confocal fluorescence microscopy images a ZEISS LSM800 with Airyscan was used. The percent cells with MDCs were quantified in each experiment at the two-hour time point. All quantifications show the mean ± standard error from three biological replicates with *n* = 100 cells per experiment. MDCs were identified as Tom70-positive, Tim50-negative structures that were enriched for Tom70 versus the mitochondrial tubule. In MDC colocalization assays, MDCs were identified as large, Tom70-enriched, spherical structures prior to assessing the co-localization of different proteins of interest.

### Fluorescence Microscopy

Fluorescence microscopy was performed as described in English et al., 2020. In brief, optical z-sections of live yeast cells were acquired with a ZEISS Axio Imager M2 equipped with a ZEISS Axiocam 506 monochromatic camera, 100x oil-immersion objective (plan apochromat, NA 1.4) or 63x oil-immersion objective (plan apochromat, NA 1.4) or a ZEISS LSM800 equipped with an Airyscan detector, 63x oil-immersion objective (plan apochromat, NA 1.4). Time-lapse fluorescence microscopy imaging was also performed as described in English et al., 2020. Briefly, overnight log-phase cultures were treated with 1 µM rapamycin for 15 minutes, harvested by centrifugation, resuspended in SD medium, and pipetted into flow chamber slides as previously described (English et al., 2020). Optical z-sections of live yeast cells were acquired with a ZEISS Airyscan LSM880 equipped with an environmental chamber set to 30°C. Widefield images were acquired with ZEN (Carl Zeiss) and processed with Fiji (Schindelin et al., 2012). Time-lapse images and super-resolution images were acquired with ZEN (Carl Zeiss) and processed using the automated Airyscan processing algorithm in ZEN (Carl Zeiss) and Fiji. Individual channels of all images were minimally adjusted in Fiji to match the fluorescence intensities between channels for better visualization. Line-scan analysis was performed on non-adjusted, single z-sections in Fiji.

### Transmission Electron Microscopy and Electron Tomography

Yeast cells were high-pressure frozen and freeze-substituted as previously described (Wilson et al., 2021). Liquid cultures of yeast cells were harvested at mid-logarithmic phase, vacuum-filtered on 0.45-μm millipore paper, loaded into 0.5-mm aluminum hats, and high pressure frozen with a Wohlwend HPF (Wohlwend, Switzerland). Cells were freeze-substituted in an Automated Freeze-Substitution machine (AFS, Leica Vienna, Austria) at −90°C in an *en bloc* preparation of 0.1% uranyl acetate and 0.25% glutaraldehyde in anhydrous acetone. Samples were then washed in pure anhydrous acetone, embedded in Lowicryl HM20 resin (Polysciences, Warrington, PA), UV polymerized at −60°C warming slowly over 4 days to room temperature (RT). The sample blocks were then stored at −20°C. These methods preserve membrane and protein structure and provide consistent *en bloc* staining for immuno-EM membrane identification (Giddings, 2003).

A Leica UC6 Ultra-Microtome was used to cut and place serial sections on Formvar-coated rhodium-plated copper slot grids (Electron Microscopy Sciences). 80 to 90-nm serial sections were cut for transmission electron microscopy (TEM) and immuno-EM experiments and 200-nm thick serial sections were cut for dual-axis tomography. For immunolabeling experiments, grids were exposed to sequential 50 µL droplets: Nonspecific antibody binding was blocked by incubation with 1% PBS + 1% dry milk (blocking solution) for 20 minutes at RT, then exposed to primary antibodies overnight at 4°C (1:500 anti-GFP) in blocking solution, washed at RT in 1% PBS with 3 sequential 50 µL drops, labeled with a secondary anti-rabbit or anti-mouse gold (depending on the primary antibody used) at RT for 1 hour (1:200 goat-anti-rabbit or goat-anti-mouse, Electron Microscopy Sciences), washed in 1% PBS with 3 sequential 50 µL drops, and finally washed in distilled water with 2 sequential 50 µL drops.

Thin cell sections were imaged with a FEI Tecnai T12 Spirit electron microscope equipped with a 120 kV LaB6 filament and AMT (2 k × 2 k) CCD. TEM of hundreds of cells per strain were used to quality control freezing, embedding, and staining. Thick sections were labeled with fiduciary 15-nm colloidal gold (British Biocell International) on both sides and tilt imaged with a Tecnai 30 (f-30, 300 kV; FEI-Company, Eindhoven, the Netherlands) with dual–tilt series images collected from +60° to −60° with 1.5° increments using a Gatan US4000 4k × 4k charge-coupled device camera (Abingdon, United Kingdom). The tilt series were imaged primarily at 19,000× magnification and repeated with a 90° rotation for dual-axis tomography (Mastronarde 1997). Tomograms were built and modeled using the IMOD software package (Kremer et al., 1996) using an iMac (Apple). MDC, mitochondria, and ER membrane models from dual-axis electron tomograms and immuno-tomograms were manually assigned from the outer leaflet every 5 nm. Immuno-gold was modeled at the same size as secondary gold (6-nm anti-mouse and 10-nm anti-rabbit, Electron Microscopy Sciences) and colored similarly to the closest membrane as indicated in the Figure Legends. Videos were made using IMOD and QuickTime Pro (Apple). Data were analyzed and graphed using Prism 9 (GraphPad).

### Protein Preparation and Immunoblotting

For western blot analysis of protein levels, yeast cultures were grown to log-phase (OD_600_= 0.5-1) and 2 OD_600_ cell equivalents were isolated by centrifugation, washed with dH_2_O and incubated in 0.1 M NaOH for five minutes at RT. Subsequently, cells were reisolated by centrifugation at 16,000 × g for ten minutes at 4°C and lysed for five minutes at 95°C in lysis buffer (10 mM Tris pH 6.8, 100 mM NaCl, 1 mM EDTA, 1 mM EGTA, 1% SDS and containing cOMPLETE protease inhibitor cocktail (Millipore Sigma)). Upon lysis, samples were denatured in Laemmli buffer (63 mM Tris pH 6.8, 2% SDS, 10% glycerol, 1 mg/ml bromophenol blue, 1% β-mercaptoethanol) for five minutes at 95°C. To separate proteins based on molecular weight, equal amounts of protein were subjected to SDS polyacrylamide gel electrophoresis and transferred to nitrocellulose membrane (Millipore Sigma) by semi-dry transfer. Nonspecific antibody binding was blocked by incubation with Tris buffered saline + 0.05% Tween-20 (TBST) containing 10% dry milk (Sigma Aldrich) for one hour at RT. After incubation with the primary antibodies at 4°C overnight, membranes were washed four times with TBST and incubated with secondary antibody (goat-anti-rabbit or donkey-anti-mouse HRP-conjugated,1:5000 in TBST + 10% dry milk, Sigma Aldrich) for 1 hour at RT. Subsequently membranes were washed twice with TBST and twice with TBS, enhanced chemiluminescence solution (Thermo Fisher) was applied and the antibody signal was detected with a BioRad Chemidoc MP system. All blots were exported as TIFFs and cropped in Adobe Photoshop CC.

### Auxin-Induced Protein Degradation

Auxin-induced protein degradation was performed essentially as described in Shetty et al., 2019 except 3-indole acetic acid (auxin) was added to a final concentration of 1 mM at the 0-time point in all experiments. All yeast strains containing the auxin-inducible degron, IAA7, were generated by endogenous C-terminal integration of yEGFP-IAA7 PCR amplified from a plasmid created for this study (Table S3) by removing 3V5 from the plasmid describe in Eng et al., 2014 by cutting with Pac1/Xba1 and replacing with yEGFP cut with similar restriction enzymes. Except Mdm12, which was C-terminally fused to AID*-6xFLAG from the constructs described in Morawska and Ulrich, 2013. Subsequently, GPD1-OsTIR1 was integrated into the *LEU2* locus, using the plasmid pNH605-pGPD1-osTIR1 digested with Swa1 as described in Chan et al., 2018. Auxin-induced protein degradation was followed by both immunoblotting from whole cell extracts and fluorescence microscopy as described above.

## QUANTIFICATION AND STATISTICAL ANALYSIS

The number of replicates, what *n* represents, and dispersion and precision measures are indicated in the Figure Legends. In general, quantifications show the mean ± standard error from three biological replicates with *n* = 100 cells per experiment. In experiments with data depicted from a single biological replicate, the experiment was repeated with the same results.

## ACKNOWLEDGEMENTS

We thank members of the A.L. Hughes and Greg Odorizzi groups for discussion and manuscript comments. We thank members of Janet. M. Shaw laboratory for providing reagents, plasmids, and early support on this project. We also thank Maya Schuldiner for gifting the yeast SWAp-Tag library described in Weill et al., 2018. Zachary Wilson, PhD, was supported by an American Heart Association Postdoctoral Fellowship, 20POST35200110, and supported by the American Cancer Society–Give Mas, Live Mas Southern Multifoods Postdoctoral Fellowship, PF-20-018-01-CCG. Research was further supported by National Institutes of Health grants T32GM007464 (A.M.E.), R35GM149202 to G.O., and GM119694 to A.L.H.

## AUTHOR CONTRIBUTIONS

Conceptualization, Z.N.W. and A.L.H.; methodology, Z.N.W, M.W., and A.M.E; formal analysis, Z.N.W. and M.W.; investigation, Z.N.W., M.W. and A.M.E.; writing – original draft, Z.N.W; writing – review and editing, Z.N.W., G.O., and A.L.H.; visualization, Z.N.W. and M.W.; supervision, G.O. and A.L.H.; funding acquisition, Z.N.W., G.O., and A.L.H.

## DECLARATION OF INTERESTS

The authors declare no competing interests.

## CONTACT FOR REAGENT AND RESOURCE SHARING

Further information and requests for resources and reagents should be directed to and will be fulfilled by the Lead Contact, Adam Hughes. All unique/stable reagents generated in this study are available from the Lead Contact without restrictions.

---

**Video 1 (related to Fig. 2). Mitochondrial-derived multilamellar structures are enriched for Tom70-GFP and exclude Tim50-mCherry.** z-series and 3D model of the tomogram shown in Figure 2.

**Video 2 (related to Fig. 3 A-C). Mitochondrial-derived compartments contain sets of paired membranes.** z-series and 3D model of the tomogram shown in Fig. 3 A-C.

**Video 3 (related to Fig. 3 D-F). Mitochondrial-derived compartments contain sets of paired membranes.** z-series and 3D model of the tomogram shown in Fig. 3 D-F.

**Video 4 (related to Fig. 4 A). Mitochondrial-derived compartments form through membrane extension intermediates.** Maximum intensity projections of yeast expressing Tom70-yEGFP and Tim50-mCherry treated with rapamycin. Images were taken every minute (min) and are shown at four frames per second.

**Video 5 (related to Fig. 4 C). Mitochondrial-derived compartments form through membrane extension intermediates.** Maximum intensity projections of yeast expressing Tom70-yEGFP treated with rapamycin. Images were taken every minute (min) and are shown at four frames per second.

## SUPPLEMENTAL FIGURE LEGENDS

**Figure S1.**
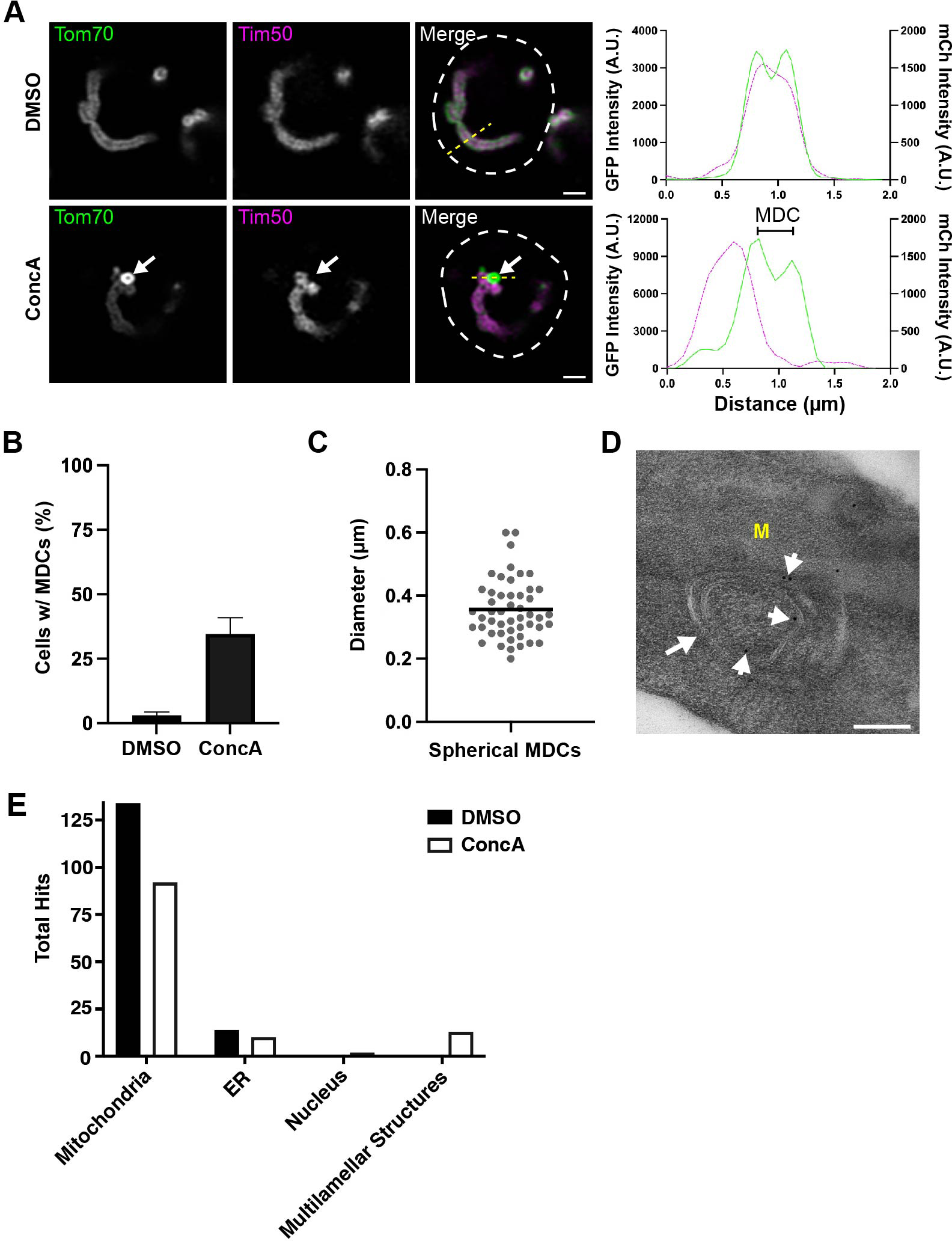
Concanamycin A (ConcA) treated yeast also produce mitochondrial-derived multilamellar structures. (A) Super-resolution confocal fluorescence microscopy images of haploid yeast expressing Tom70-yEGFP and Tim50-mCherry treated with either DMSO or 500 nM ConcA. MDCs are indicated by white arrows. Scale Bar= 1 µm. Yellow line marks the position of the line-scan fluorescence intensity profile shown to the right. Left and right Y axis correspond to Tom70-yEGFP and Tim50-mCherry fluorescence intensity, respectively. Bracket marks MDC. (B) Quantification of MDC formation in DMSO or ConcA treated yeast. Error bars show mean ± standard error of three replicates, *n* ≥ 100 cells per replicate. (C) Scatter plot showing the diameter of ConcA-induced MDCs. Black line indicates the mean (0.36 µm) of *n* = 49 MDCs. (D) Thin-section TEM analysis of 80-nm cell sections from the same yeast strain analyzed above treated with 500 nM ConcA. Sections were stained with monoclonal antibodies targeting GFP and secondary antibodies conjugated to 10-nm gold particles. White arrow: multilamellar structures, White arrowheads point to gold particles, M: mitochondria. Scale Bar= 200 nm. (E) Quantification of the total anti-GFP immunogold particles that labeled the indicated cell structures from an analysis of >100 cell-sections from yeast expressing Tom70-yEGFP that were treated with either DMSO (Vehicle) or 500 nM ConcA.

**Figure S2.**
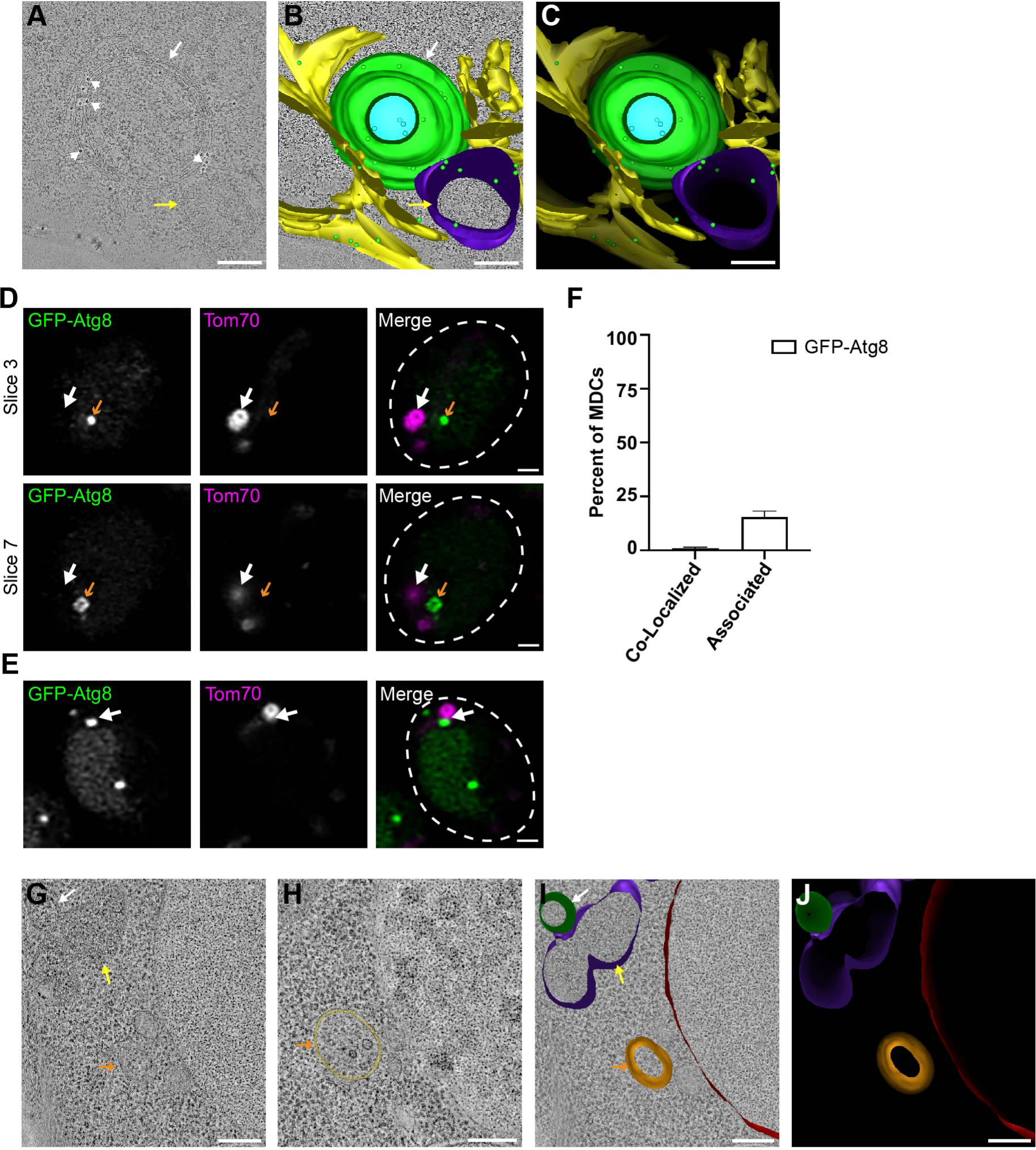
MDCs are not bound by autophagosomal membranes. (A - C) 2D cross sections and a 3D model derived from 5 serial 90-nm cell sections of the same yeast strain analyzed in Fig. 1. Sections were immunolabeled with antibodies targeting GFP and secondary antibodies conjugated to 10-nm gold particles. Scale bar= 200 nm. See associated Video S1. (A) Tomograph of a large, multilamellar structure strongly labeled with antibodies targeting Tom70-yEGFP. Yellow arrow: mitochondria, white arrow: multilamellar structure, white arrowheads point to gold particles. (B) Model overlay of the tomograph shown in A. (C) 3D model of a large, multilamellar structure bound by two closely-apposed paired membranes. The limiting membrane of the outer doublet membrane is labeled green, while the internal doublet membrane is labeled cyan. Mitochondria: purple, ER: yellow, and 10-nm gold particles are overlayed with green spheres. (D and E) Representative super-resolution confocal fluorescence microscopy images of yeast expressing Tom70-mCherry and sfGFP-Atg8 after a 2-hour treatment with 200 nM rapamycin. White arrow marks MDC, orange arrow marks sfGFP-Atg8 positive structures. The images shown in D are from the same cell but show different z-slices, which are labeled. (F) Quantification of the frequency sfGFP-Atg8 foci were co-localized or closely associated to Tom70-mCherry-marked MDCs. Error bars show mean ± standard error of three replicates, *n* ≥ 100 MDCs per replicate. (G-J) 2D cross sections and 3D models from different views of the same tomogram shown in Fig. 3 A-C. Scale bars= 200 nm. (G and H) Two different 2D cross sections from a larger view of the same tomogram shown in Fig. 3 A-C. White arrow: MDC, yellow arrow: mitochondria, orange arrow: autophagosome. (I) Model overlay of the tomograph shown in G. (J) 3D model of a vacuole-associated autophagosome labeled in orange, MDC labeled in green, mitochondria labeled purple and the vacuole is labeled in red.

**Figure S3.**
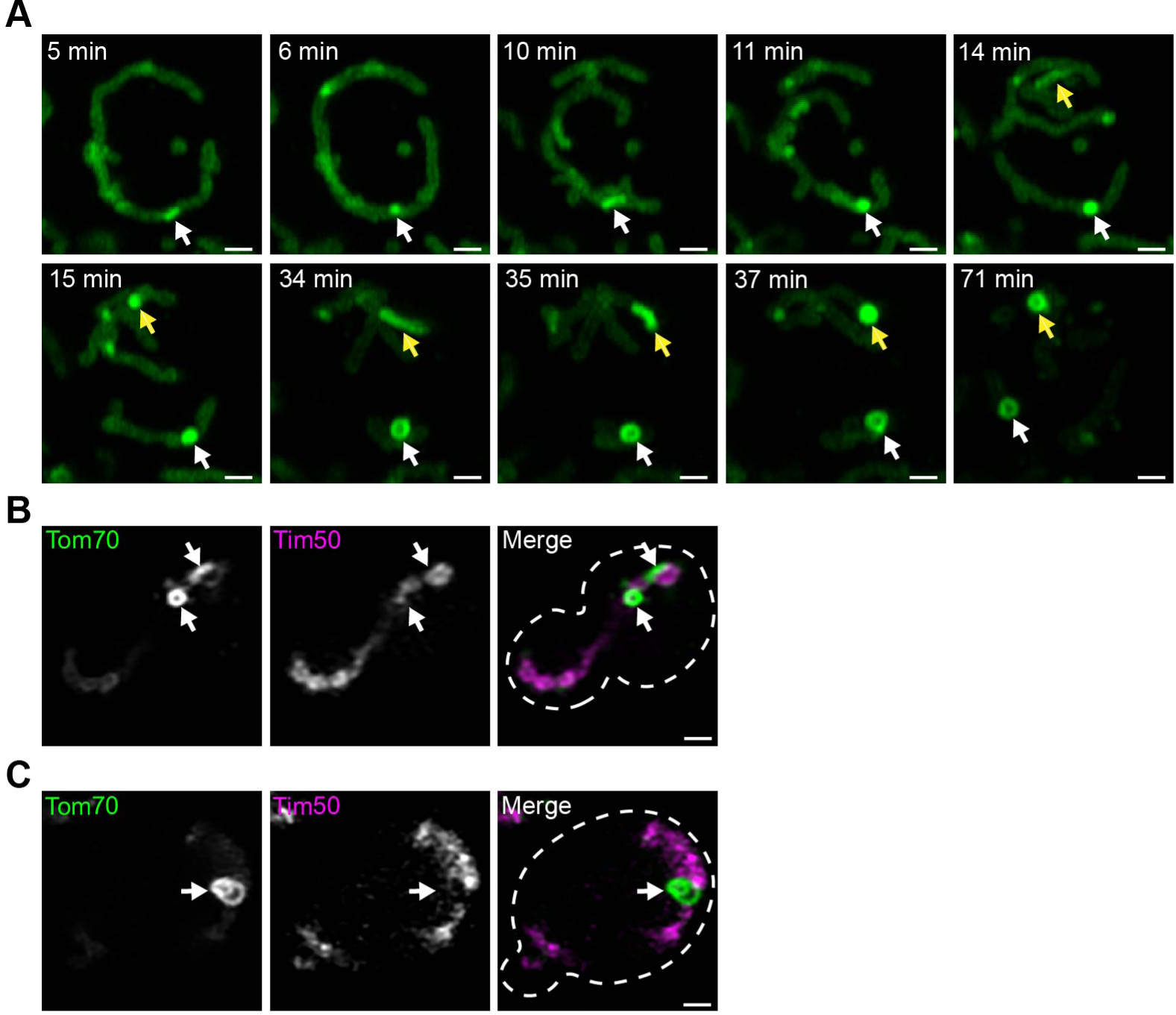
Mitochondrial-derived compartments form through membrane extension intermediates. (A) Super-resolution time-lapse images of rapamycin-induced MDC formation in yeast cells expressing Tom70-yEGFP. Images were acquired over 120 minutes (min). White and yellow arrows mark two different MDCs. Scale bar = 1 µm. See associated Video S2. (B) Representative super-resolution confocal fluorescence microscopy image of a cell with two MDCs with different morphologies. MDCs are marked by white arrows. (C) Representative super-resolution confocal fluorescence microscopy image of an MDC with resolvable layers of Tom70-yEGFP. MDC is marked by the white arrow.

**Video S1 (related to Fig. S2). Mitochondrial-derived compartments form through membrane extension intermediates.** z-series and 3D model of the tomogram shown in Fig. S2 A-C.

**Video S2 (related to Fig. S3 A). Mitochondrial-derived compartments form through membrane extension intermediates.** Maximum intensity projections of yeast expressing Tom70-yEGFP treated with rapamycin. Images were taken every minute (min) and are shown at four frames per second.

## Notes

### Competing Interest Statement

The authors have declared no competing interest.

