## Supplementary material for "Mitochondrial-Derived Compartments are Multilamellar Domains that Encase Membrane Cargo and Cytosol": Table S3

**Table S3. Bacterial strains, chemicals, plasmids, and software used in this study**

| **Reagent or resource** | | **Source** | | **Identifier** |
| --- | --- | --- | --- | --- |
| **Antibodies** | | | | |
| Mouse monoclonal anti-GFP clones 7.1 and 13.1 (dilution 1:1000 or 1:200 immuno-EM) | Roche | | Cat # 11814460001; RRID:AB_390913 | |
| Mouse monoclonal anti-PGK1 clone 22C5D8 (dilution 1:5000) | Abcam | | Cat # ab113687; RRID:AB_10861977 | |
| Mouse monoclonal anti-Por1 clone 16G9E6BC4 (dilution 1:1000) | Abcam | | RRID:AB_10865182; Cat # ab110326 | |
| Rabbit anti-mCherry (dilution 1:200 immuno-EM) | Pearson Lab | | N/A | |
| Rabbit polyclonal anti-Aco1 (dilution 1:2000) | Shaw Lab | | N/A | |
| Rabbit polyclonal anti-Aac2 (dilution 1:5000) | Shaw Lab | | N/A | |
| Rabbit polyclonal anti-Om45 (dilution 1:2000) | Shaw Lab | | N/A | |
| Rabbit polyclonal anti-Tom20 (dilution 1:1000) | Pfanner Lab | | N/A | |
| Rabbit polyclonal anti-Tom71 (dilution 1:1000) | Pfanner Lab | | N/A | |
| Goat-anti-Mouse IgG/IgM 6 nm, Goat-anti-Rabbit 10 nm Kit (dilution 1:250) | Electron Microscopy Sciences | | Cat # 25544 | |
| **Bacterial strains** | | | | |
| *Escherichia coli* DH5α | | N/A | | N/A |
| **Chemicals, peptides, and recombinant proteins** | | | | |
| 3-Indoleacetic acid (Auxin) | | Sigma-Aldrich | | Cat # I3750 ; CAS # 87-51-4‎ |
| Casamino acids | | US Biological | | Cat # 0012501A; CAS # 65072-00-6 |
| Concanamycin A | | Santa Cruz Biotechnology | | Cat # sc-202111; CAS # 80890-47-7 |
| Concanavalin A | | Sigma-Aldrich | | Cat # L7647; CAS # 11028-71-0 |
| Dimethyl sulfoxide (DMSO) | | Sigma-Aldrich | | Cat # D2650; CAS # 67-68-5 |
| DTT | | Gold Biotechnology | | Cat # DTT10; CAS # 27565-41-9 / 3483-12-3 |
| Lyticase | | Sigma-Aldrich | | Cat # L2524; CAS #  37340-57-1 |
| Rapamycin | | LC Laboratories | | Cat # R-5000; CAS # 53123-88-9 |
| Fiduciary Gold, 15nm | | Aurion | | Cat # 415.011 |
| Critical commercial assays | | | | |
| Bicinchoninic Acid Protein Assay | | Thermo Fisher | | Cat # 23227 |
| **Plasmids** | | | | |
| Plasmid: pRS315-FZO1 | | Hermann et al., 1998  (Shaw Lab) | | B1293 |
| Plasmid: pRS415-*fzo1-1* | | Hermann et al., 1998  (Shaw Lab) | | B1016 |
| Plasmid: pFA6-GFP-IAA7::URA3 | | This Study | | B3813 |
| Plasmid: pHYG-AID*-6FLAG | | Morawska and Ulrich, 2013 Addgene | | Plasmid # 99519 |
| Plasmid: pKT127 | | (Sheff and Thorn, 2004); Addgene | | Plasmid # 8728 |
| Plasmid: pKT127-mCherry | | Daniel Gottschling (Calico) | | N/A |
| Plasmid: pKT128 | | Sheff and Thorn, 2004; Addgene | | Plasmid # 8729 |
| Plasmid: pRS305 | | Sikorski and Hieter, 1989 | | N/A |
| Plasmid: pRS306 | | Sikorski and Hieter, 1989 | | N/A |
| Plasmid: pRS40Hyg | | Daniel Gottschling (Calico) | | N/A |
| **Software and algorithms** | | | | |
| IMOD | | Kremer et al. 1996 | | Version 4.11 |
| FIJI | | Schindelin *et al*., 2012 | | Version 1 |
| Prism | | GraphPad Software, Inc. | | Version 9 |
| SnapGene | | GSL Biotech | | Version 4.2 |
| ZEN Blue Edition | | Carl Zeiss Microscopy | | Version 2.6 |
